# Germline status and micronutrient availability regulate a somatic mitochondrial quality control pathway via short-chain fatty acid metabolism

**DOI:** 10.1101/2024.05.20.594820

**Authors:** James P. Held, Nadir H. Dbouk, Adrianna M. Strozak, Lantana K. Grub, Hayeon Ryou, Samantha H. Schaffner, Maulik R. Patel

**Affiliations:** Department of Biological Sciences, Vanderbilt University, Nashville, TN, USA; Department of Cell & Developmental Biology, Vanderbilt University School of Medicine, Nashville, TN, USA; Evolutionary Studies, Vanderbilt University, VU Box #34-1634, Nashville, TN, USA; Diabetes Research and Training Center, Vanderbilt University School of Medicine, Nashville, TN, USA; Quantitative Systems Biology Center, Vanderbilt University, Nashville, TN, USA

## Abstract

Reproductive status, such as pregnancy and menopause in women, profoundly influences metabolism of the body. Mitochondria likely orchestrate many of these metabolic changes. However, the influence of reproductive status on somatic mitochondria and the underlying mechanisms remain largely unexplored. We demonstrate that reproductive signals modulate mitochondria in the *Caenorhabditis elegans* soma. We show that the germline acts via an RNA endonuclease, HOE-1, which despite its housekeeping role in tRNA maturation, selectively regulates the mitochondrial unfolded protein response (UPR^mt^). Mechanistically, we uncover a fatty acid metabolism pathway acting upstream of HOE-1 to convey germline status. Furthermore, we link vitamin B12’s dietary intake to the germline’s regulatory impact on HOE-1-driven UPR^mt^. Combined, our study uncovers a germline-somatic mitochondrial connection, reveals the underlying mechanism, and highlights the importance of micronutrients in modulating this connection. Our findings provide insights into the interplay between reproductive biology and metabolic regulation.

## MAIN

Reproduction is a fundamental biological process. Considering the substantial energetic investment required for reproduction, organisms display acute sensitivity to various stages of their reproductive phases and instigate metabolic changes to accommodate these demands ^1–3^. As cellular hubs for metabolism, mitochondria likely lie at the heart of many of these metabolic adjustments ^4–6^. However, the impact of reproductive status on mitochondria within somatic cells and the molecular mechanisms that underlie these changes remain largely unexplored.

To elucidate the regulatory role that the germline exerts on somatic mitochondria, we focused on the mitochondrial unfolded protein response (UPR^mt^) ^7,8^. Best characterized in *Caenorhabditis elegans*, UPR^mt^ is a key mitochondrial quality control pathway activated in response to a variety of mitochondrial perturbations ^9^. UPR^mt^ entails expression of numerous target genes including mitochondrial proteases that degrade damaged or improperly folded proteins, and chaperones that increase the protein folding capacity of mitochondria to accommodate newly synthesized proteins ^10^. UPR^mt^ also boosts the expression of mitochondrial protein transport machinery, aiding in mitochondrial import recovery. Finally, activation of UPR^mt^ results in metabolic remodeling and tolerance to reactive oxygen species toxicity. Taken together, as a vital mechanism for mitochondrial quality control that reconfigures mitochondrial function, UPR^mt^ is optimally situated to adapt mitochondrial responses to alterations in reproductive status.

We recently reported a surprising function of the *C. elegans* homolog of ELAC2, known as HOE-1, as a central regulator of UPR^mt^ ^11^. ELAC2, also referred to as RNaseZ, is an RNA endonuclease that cleaves 3’-trailer sequences from nascent tRNAs in both the nucleus and mitochondria ^12–15^. This cleavage is a crucial step in tRNA maturation—essential for subsequent modifications and amino acid charging. Mutations in ELAC2 are associated with prostate cancer and are known to cause hypertrophic cardiomyopathy ^15–22^. Notably, our investigations revealed that it is the nuclear isoform of HOE-1, rather than its mitochondrial counterpart, that governs UPR^mt^ dynamics. Preventing HOE-1 nuclear import resulted in the attenuation of UPR^mt^, while increasing HOE-1 nuclear levels through the deletion of its nuclear export signal induced a robust UPR^mt^ response that depended on ATFS-1, the central transcription factor necessary for UPR^mt^ ^23^. Furthermore, either introducing a mutation disrupting HOE-1’s enzymatic function or inhibiting the canonical tRNA exportin effectively suppressed HOE-1-triggered UPR^mt^, suggesting that HOE-1 exerts its regulatory influence via its tRNA or tRNA-like substrates. Consistent with its involvement in UPR^mt^, the nuclear localization of HOE-1 is dynamically responsive to ATFS-1 activity. Taken together, there is an unusual connection between UPR^mt^, a response specific to mitochondria, and HOE-1, an RNA processing enzyme primarily associated with an essential housekeeping function.

Here, we report our discovery that the germline exerts non-cell autonomous control over HOE-1 triggered UPR^mt^ in the soma. Moreover, HOE-1 activates UPR^mt^ specifically in the intestine— functionally analogous to metabolic organs in mammals, and only after animals have reached reproductive adulthood. The germline exerts its effects by promoting nuclear accumulation of HOE-1. Mechanistically, we show that a mitochondrially localized propionate shunt pathway mediated by an acyl-CoA dehydrogenase, ACDH-1, regulates HOE-1 in a germline cell non-autonomous manner. Specifically, we isolated gain-of-function and loss-of-function mutations in key pathway enzymes including in ACDH-1 that, combined with the genetic analysis of other enzymes, identified the specific metabolic step regulated by the germline to control HOE-1. Interestingly, the expression of ACDH-1 and other enzymes in the propionate metabolism pathway is sensitive to vitamin B12 availability. Consequently, we find that the germline regulation of HOE-1 triggered UPR^mt^ is completely dependent on the dietary presence of vitamin B12. In summary, our findings unveil an interplay where germline status and micronutrient availability emerge as crucial determinants in the orchestration of HOE-1-triggered UPR^mt^.

## RESULTS

### Increased nuclear activity of HOE-1 has a strong mitochondrial signature

We previously demonstrated that enhancing the nuclear localization of HOE-1 by disrupting its nuclear export signal (allele mpt67, denoted as *hoe-1*(ΔNES)), robustly activates UPR^mt^ while showing no activation of the endoplasmic reticulum UPR ^11^. This suggests specific involvement of HOE-1 in mitochondrial regulation. To comprehensively assess the broader cellular consequences of increased nuclear activity of HOE-1, we conducted RNA sequencing on wildtype vs *hoe-1*(ΔNES) adult animals, which revealed a unique transcriptional profile (Figures S1A and S1B). This was conducted in a germline-less *glp-1* temperature sensitive background to specifically assess the somatic consequences. Our analysis revealed 1,110 significantly upregulated genes and 838 significantly downregulated genes in *hoe-1*(ΔNES) animals, with an adjusted p-value <0.05 and a log_2_ fold change >1 or <-1, respectively (Figure 1A). Notably, gene ontology (GO) analysis of significantly upregulated genes showed a significant overrepresentation of mitochondrial associated genes (Figures 1B). Knockdown of ATFS-1 strongly compromised the differentially expressed gene profile (Figures 1C). Additionally, high confidence *atfs-1* target genes ^24^, *clec-17*, *cyp-14A4*, *hsp-6*, *hsp-60*, *hrg-9*, and *K09E9.1*, exhibited upregulation in *hoe-1*(ΔNES) animals, and this effect was completely abolished on *atfs-1* RNAi (Figures 1D). Taken together, these data demonstrate an outsized role of HOE-1 in regulating the expression of genes associated with mitochondrial function, many through UPR^mt^.

**Figure 1:**
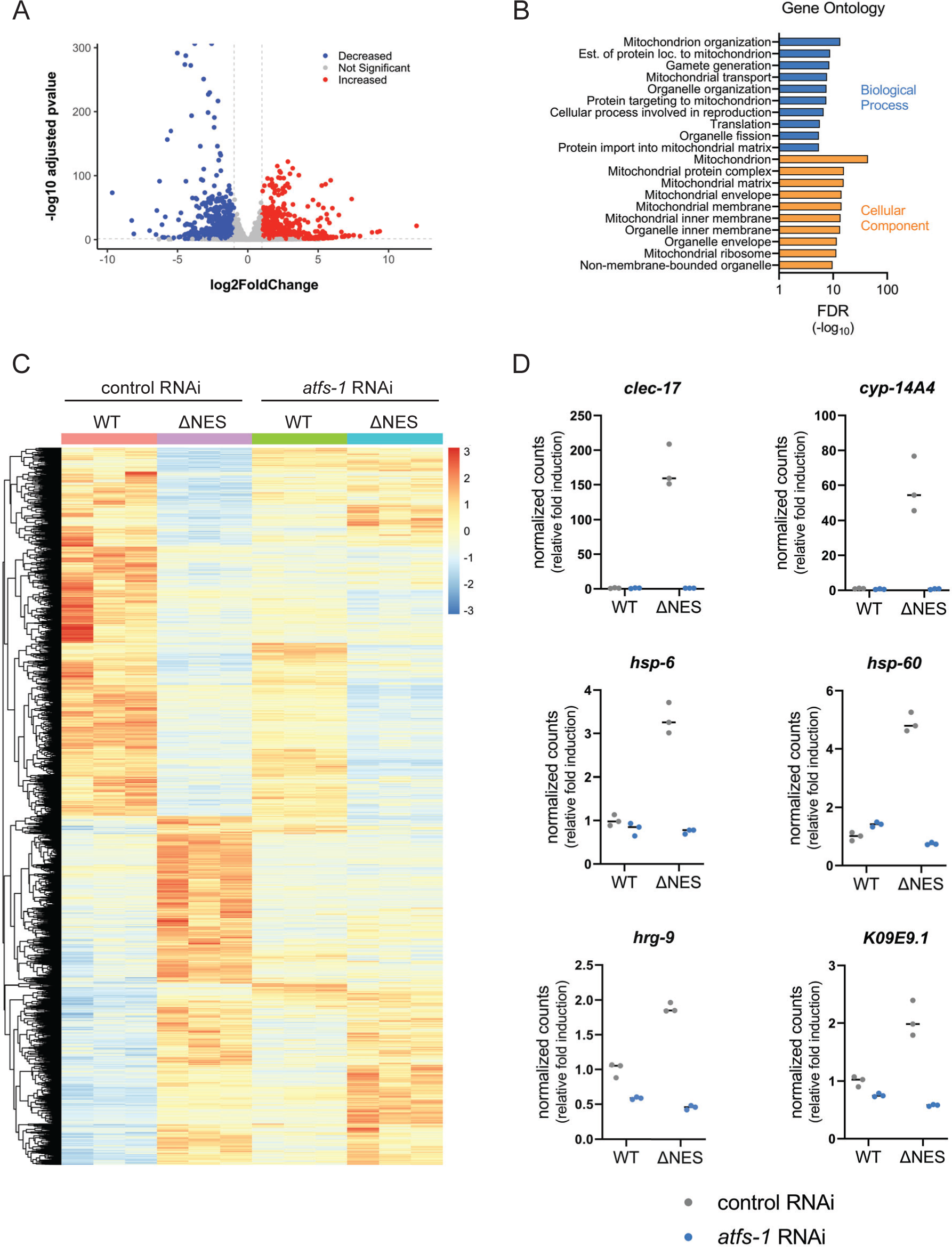
Increased nuclear activity of HOE-1 has a strong mitochondrial signature. (A) A volcano plot of gene expression differences between wildtype and *hoe-1*(ΔNES) animals on control RNAi. A log_2_ fold change cutoff of 1 or -1 (vertical dashed gray line) and a -log_10_ adjusted p-value cutoff of p < 0.05 (horizonal dashed gray line) are included. Genes with no significant change, genes with a significant increase in gene expression and genes with significantly decreased expression are represented by gray, red and blue points respectively. (B) Gene ontology analysis of genes significantly upregulated with a log_2_ fold change >1 in *hoe-1*(ΔNES) animals relative to wildtype. Shown are the 10 most significant terms for each analysis by false discovery rate (FDR). (C) Heat map of gene expression in wildtype (WT) and *hoe-1*(ΔNES) animals on control and *atfs-1* RNAi. Gene expression is visualized by a z-score calculated on a gene-by-gene basis. A z-score equal to zero is yellow, positive z-scores are red, and negative z-scores are blue. (D) Normalized transcript counts from RNA-seq of high-confidence *atfs-1* target genes *clec-17*, *cyp-14A4*, *hsp-6*, *hsp-60*, *hrg-9*, and *K09E9.1* in wildtype (WT) and *hoe-1*(ΔNES) animals on control and *atfs-1* RNAi.

### HOE-1-dependent UPR^mt^ has distinct features

In characterizing HOE-1 dependent UPR^mt^, we identified multiple features that differ between HOE-1 UPR^mt^ and UPR^mt^ induced by directly inducing mitochondrial stress. A loss-of-function mutation in the electron transport chain subunit *nuo-6* (allele qm200, denoted as *nuo-6-/-*)^25^ activates UPR^mt^ across somatic tissues, as evidenced by the *hsp-6*p::GFP reporter activation (Figures 2A and 2B). This is consistent with previous reports that systemic mitochondrial stress or constitutive UPR^mt^ activation by *atfs-1* gain-of-function mutants induce pan-somatic UPR^mt^ ^26,27^. In contrast, *hoe-1*(ΔNES) induces the UPR^mt^ reporter specifically in the intestine (Figures 2A and 2B). Additionally, we note that *nuo-6-/-* animals initiate UPR^mt^ early in development, persisting into adulthood (Figures 2C, 2D, and S2A). In contrast, UPR^mt^ activation in *hoe-1*(ΔNES) animals is delayed until late in development, achieving robust activation only in adulthood (Figures 2C, 2D, and S2A).

**Figure 2:**
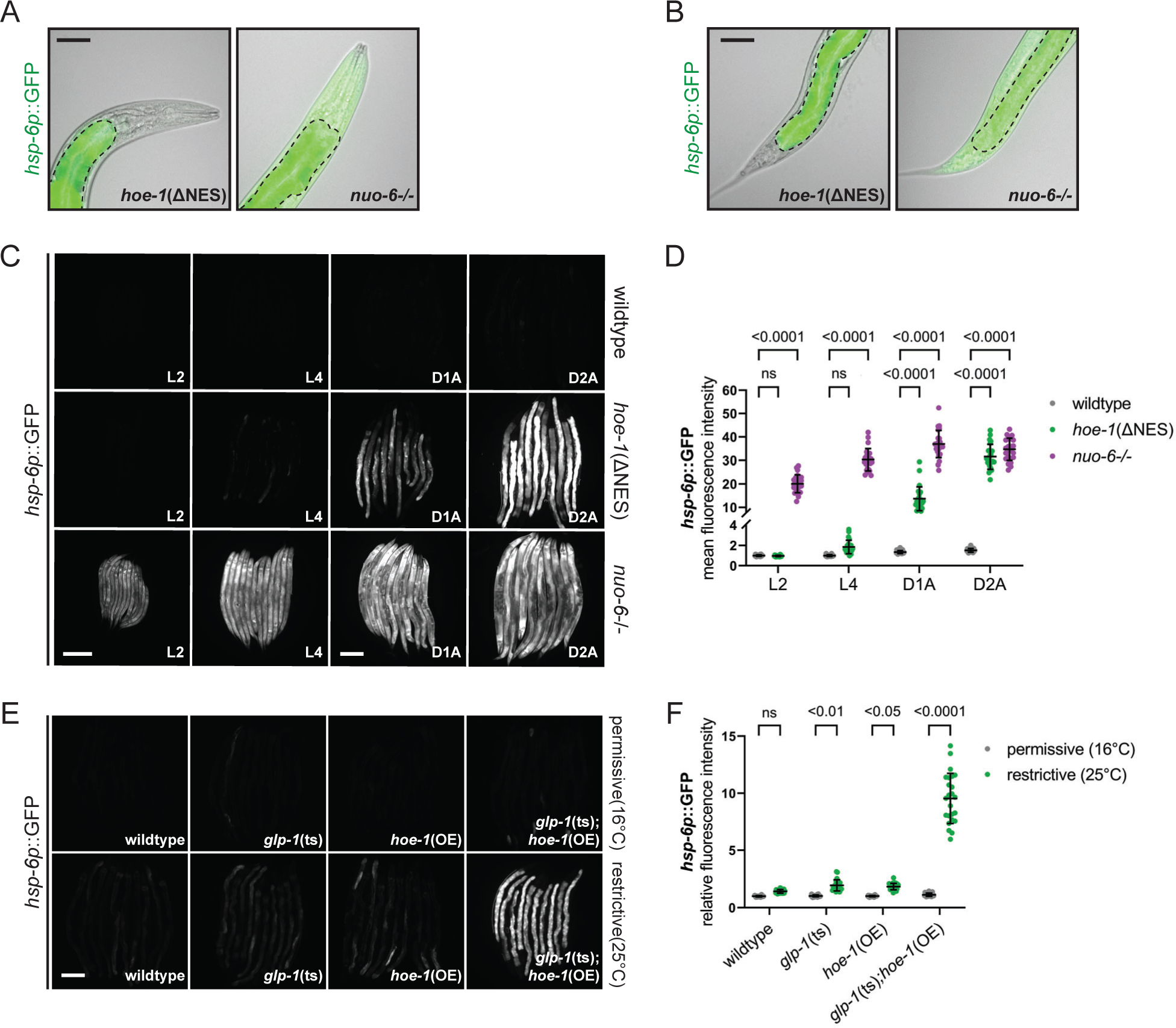
HOE-1-dependent UPR^mt^ has distinct features. Fluorescence overlayed brightfield image of UPR^mt^ reporter (*hsp-6p*::GFP) in the (A) anterior and (B) posterior of *hoe-1*(ΔNES), and *nuo-6* loss-of-function (*nuo-6-/-*) animals. The intestine is traced with a dotted line. Scale bar 50 μm. (C) Fluorescence images of UPR^mt^ reporter (*hsp-6p*::GFP) activation in wildtype, *hoe-1*(ΔNES), and *nuo-6-/-* animals across development: larval stage 2 (L2), larval stage 4 (L4), day 1 adult (D1A), day 2 adult (D2A). Scale bar 200 μm – L2 and L4 share scale bar, D1A and D2A share scale bar. (D) Relative mean fluorescence intensity quantification of *hsp-6p*::GFP in wildtype, *hoe-1*(ΔNES), and *nuo-6-/-* animals across development normalized to wildtype L4 animals (n=24, mean and SD shown, ordinary two-way ANOVA with Tukey’s multiple comparisons test). (E) Fluorescence images of UPR^mt^ reporter (*hsp-6p*::GFP) activation in day 2 adult wildtype, *glp-1* temperature-sensitive loss-of-function (*glp-1*(ts)), *hoe-1* somatic overexpression (*hoe-1*(OE)), and *glp-1*(ts);*hoe-1*(OE) animals at both a permissive (16°C – *glp-1*(ts) are fertile) and restrictive (25°C – *glp-1*(ts) are sterile) temperature. Scale bar 200 μm. (F) Relative mean fluorescence intensity quantification of *hsp-6p*::GFP in wildtype, *glp-1*(ts), *hoe-1*(OE), and *glp-1*(ts);*hoe-1*(OE) animals at both a permissive (16°C) and restrictive (25°C) temperature (n=24, mean and SD shown, ordinary two-way ANOVA with Tukey’s multiple comparisons test).

Given these distinctive features of HOE-1-dependent UPR^mt^, we aimed to identify contexts that recapitulate these outcomes. One major physiological change that coincides with the transition to adulthood in *C. elegans* is the proliferation of the germline ^28^. Thus, we hypothesized that germline status may contribute to HOE-1-dependent UPR^mt^. Indeed, we find that compromising the germline (via conditional *glp-1* loss-of-function allele e2144, denoted *glp-1*(ts)) or pan-somatically overexpressing HOE-1 (single copy knock-in strain mptSi1, denoted as *hoe-1*(OE)) individually only mildly activates UPR^mt^, but their combined effect robustly activates UPR^mt^, akin to *hoe-1*(ΔNES) animals (Figures 2E, 2F, and S2B). These findings suggest that germline status can cell non-autonomously regulate HOE-1-induced UPR^mt^.

Additionally, *hoe-1*(OE); *hoe-1* loss-of-function (allele mpt31, denoted as *hoe-1*-/-) double mutant animals develop normally into adults but have germline defects (since *hoe-1*(OE) only rescues somatic defects of *hoe-1*-/-animals) (Figure S2C). These animals exhibit robust activation of UPR^mt^ post-developmentally like that in *hoe-1*(ΔNES) and *glp-1*(ts);*hoe-1*(OE) animals (Figures S2D, S2E and S2F). These findings are consistent with the *glp-1*(ts) experiment, suggesting that overexpression of HOE-1 is sufficient to induce UPR^mt^ but only when reproduction is compromised. Taken together, our findings reveal that the germline exerts cell non-autonomous control over HOE-1 dependent UPR^mt^.

### Forward genetic screen for HOE-1-like UPR^mt^ activators yields *acdh-1* gain-of-function mutants

To unravel how the germline regulates HOE-1 dependent UPR^mt^ in the soma, we conducted a forward genetic mutagenesis screen of UPR^mt^ reporter animals, to identify genes with UPR^mt^ activation characteristics similar to HOE-1. We isolated mutants where UPR^mt^ is activated post-developmentally and specifically in the intestine (Figure 3A). We recovered six independent mutants from this screen (Figures 3B and S3A). Interestingly, in two of the mutant strains, whole genome sequencing revealed missense mutations in the *acdh-1* gene that encodes an acyl-CoA dehydrogenase (mpt134: G294E and mpt136: A365V) (Figure 3C). CRISPR-mediated introduction of both mutations independently in a clean genetic background activated UPR^mt^ post-developmentally and specifically in the intestine, confirming their causality (Figures 3D–F, S3B, and S3C).

**Figure 3:**
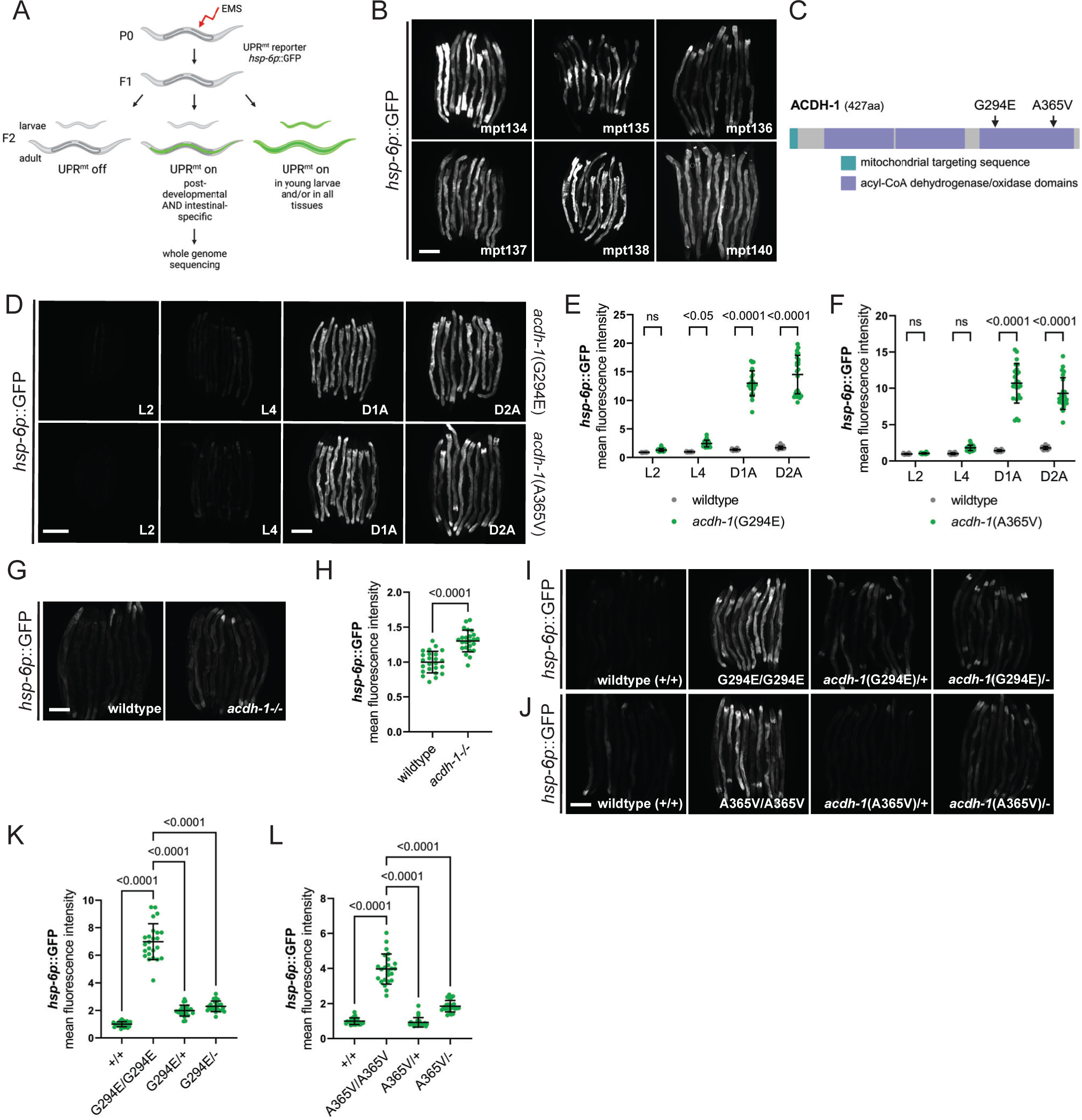
Forward genetic screen reveals gain-of-function acdh-1 mutants induce post-developmental, intestinal specific UPR^mt^. (A) Schematic outline of forward genetic mutagenesis screen. (B) Fluorescence images of UPR^mt^ reporter (*hsp-6p*::GFP) activation in mutant strains recovered from forward genetic screen. Animals imaged as day 2 adults. Scale bar 200 μm. (C) Protein schematic of ACDH-1. Missense mutations identified in mutant strain mpt134 (G294E) and mpt136 (A365V) are indicated. (D) Fluorescence images of UPR^mt^ reporter (*hsp-6p*::GFP) activation in *acdh-1*(G294E) and *acdh-1*(A365V) animals across development: larval stage 2 (L2), larval stage 4 (L4), day 1 adult (D1A), day 2 adult (D2A). Scale bar 200 μm – L2 and L4 share scale bar, D1A and D2A share scale bar. (E) Relative mean fluorescence intensity quantification of *hsp-6p*::GFP in wildtype and *acdh-1*(G294E) animals across development normalized to wildtype L4 animals (n=24, mean and SD shown, ordinary two-way ANOVA with Tukey’s multiple comparisons test). (F) Relative mean fluorescence intensity quantification of *hsp-6p*::GFP in wildtype and *acdh-1*(A365V) animals across development normalized to wildtype L4 animals (n=24, mean and SD shown, ordinary two-way ANOVA with Tukey’s multiple comparisons test). (G) Fluorescence images of UPR^mt^ reporter (*hsp-6p*::GFP) activation in day 2 adult wildtype and *acdh-1* loss-of-function (*acdh-1-/-*) animals. Scale bar 200 μm (H) Relative mean fluorescence intensity quantification of *hsp-6p*::GFP in day 2 adult wildtype and *acdh-1-/-* animals (n=24, mean and SD shown, unpaired t-test). (I) Fluorescence images of UPR^mt^ reporter (*hsp-6p*::GFP) activation in day 2 adult wildtype (+/+), *acdh-1*(G294E) (G294E/G294E), *acdh-1*(G294E) x wildtype trans-heterozygous (G294E/+), and *acdh-1*(G294E) x *acdh-1* loss-of-function trans-heterozygous (G294E/-) animals. Scale bar 200 μm. (J) Fluorescence images of UPR^mt^ reporter (*hsp-6p*::GFP) activation in day 2 adult wildtype (+/+), *acdh-1*(A365V) (A365V/A365V), *acdh-1*(A365V) x wildtype trans-heterozygous (A365V/+), and *acdh-1*(A365V) x *acdh-1* loss-of-function trans-heterozygous (A365V/-) animals. Scale bar 200 μm. (K) Relative mean fluorescence intensity quantification of *hsp-6p*::GFP in day 2 adult wildtype (+/+), *acdh-1*(G294E) (G294E/G294E), *acdh-1*(G294E) x wildtype trans-heterozygous (G294E/+), and *acdh-1*(G294E) x *acdh-1* loss-of-function trans-heterozygous (G294E/-) animals (n=24, mean and SD shown, ordinary one-way ANOVA with Tukey’s multiple comparisons test). (L) Relative mean fluorescence intensity quantification of *hsp-6p*::GFP in day 2 adult wildtype (+/+), *acdh-1*(A365V) (A365V/A365V), *acdh-1*(A365V) x wildtype trans-heterozygous (A365V/+), and *acdh-1*(A365V) x *acdh-1* loss-of-function trans-heterozygous (A365V/-) animals (n=24, mean and SD shown, ordinary one-way ANOVA with Tukey’s multiple comparisons test).

To ascertain the functional nature of these mutations, we initially examined whether a known loss-of-function mutation in *acdh-1* (allele ok1489, denoted as *acdh-1*-/-), could activate UPR^mt^. In contrast to the *acdh-1* point mutants, loss of *acdh-1* only mildly activates the UPR^mt^ reporter suggesting that *acdh-1*(G294E) and *acdh-1*(A365V) are unlikely loss-of-function (Figures 3G, 3H and S3D). To further examine the characteristics of the *acdh-1* missense mutations, we produced trans-heterozygous animals by crossing *acdh-1*(G294E) and *acdh-1*(A365V) with both wildtype (+) and loss-of-function (-) alleles of *acdh-1*. Notably, only homozygous G294E and A365V animals exhibit robust UPR^mt^ activation, whereas trans-heterozygous combinations (G294E/+, G294E/-, A365V/+, and A365V/-) display little to no UPR^mt^ reporter activation (Figures 3I–L and S3E). These results confirm that both *acdh-1* missense mutations are gain-of-function that do not act dominantly. In conclusion, increased activity of ACDH-1 induces UPR^mt^ in a manner analogous to HOE-1.

### ACDH-1 triggered UPR^mt^ is sensitive to vitamin B12 levels

ACDH-1 is known to play an important role in propionate metabolism ^29^. Propionyl-CoA, a three-carbon fatty acid, is generated through the breakdown of certain amino acids and the final stage of odd-chain fatty acid β-oxidation. In contrast to even-chain fatty acids, which yield two-carbon acetyl-CoA molecules during β-oxidation, propionyl-CoA undergoes additional metabolic transformations (Figure 4A) ^30^. Specifically, a series of enzymatic reactions converts propionyl-CoA into succinyl-CoA. Notably, one of these enzymes, methylmalonyl-CoA mutase, relies on vitamin B12 as a cofactor ^31,32^. However, when vitamin B12 is deficient, propionyl-CoA takes an alternative route, breaking down into acetyl-CoA ^33^. Importantly, ACDH-1 initiates the first step of the vitamin B12 independent pathway, and its expression is regulated by vitamin B12 availability ^29,33^. *Acdh-1* exhibits high expression under the standard laboratory diet of *E. coli* strain OP50, which is low in vitamin B12 ^33,34^. Conversely, vitamin B12 supplementation inhibits *acdh-1* expression^29,33^. Thus, we hypothesized that if *acdh-1*(G294E) and *acdh-1*(A365V) activate UPR^mt^ via increased *acdh-1* activity, then vitamin B12 supplementation should counteract UPR^mt^ activation. Indeed, vitamin B12 supplementation reduces UPR^mt^ reporter activation to wildtype levels in both *acdh-1* mutants (Figures 4B, 4C, S4A, and S4B). Moreover, UPR^mt^ reporter activation was completely abolished by vitamin B12 supplementation in both mutant strains that we recovered from the screen containing mutations in *acdh-1* (Figures S4C–S4F). This vitamin B12 dependence is specific to ACDH-1-induced UPR^mt^, as vitamin B12 supplementation has minimal effect on UPR^mt^ in *nuo-6-/-* animals (Figures 4D, 4E, and S4G). Taken together, these findings illustrate that ACDH-1 activates UPR^mt^ in a manner sensitive to the dietary levels of vitamin B12.

**Figure 4:**
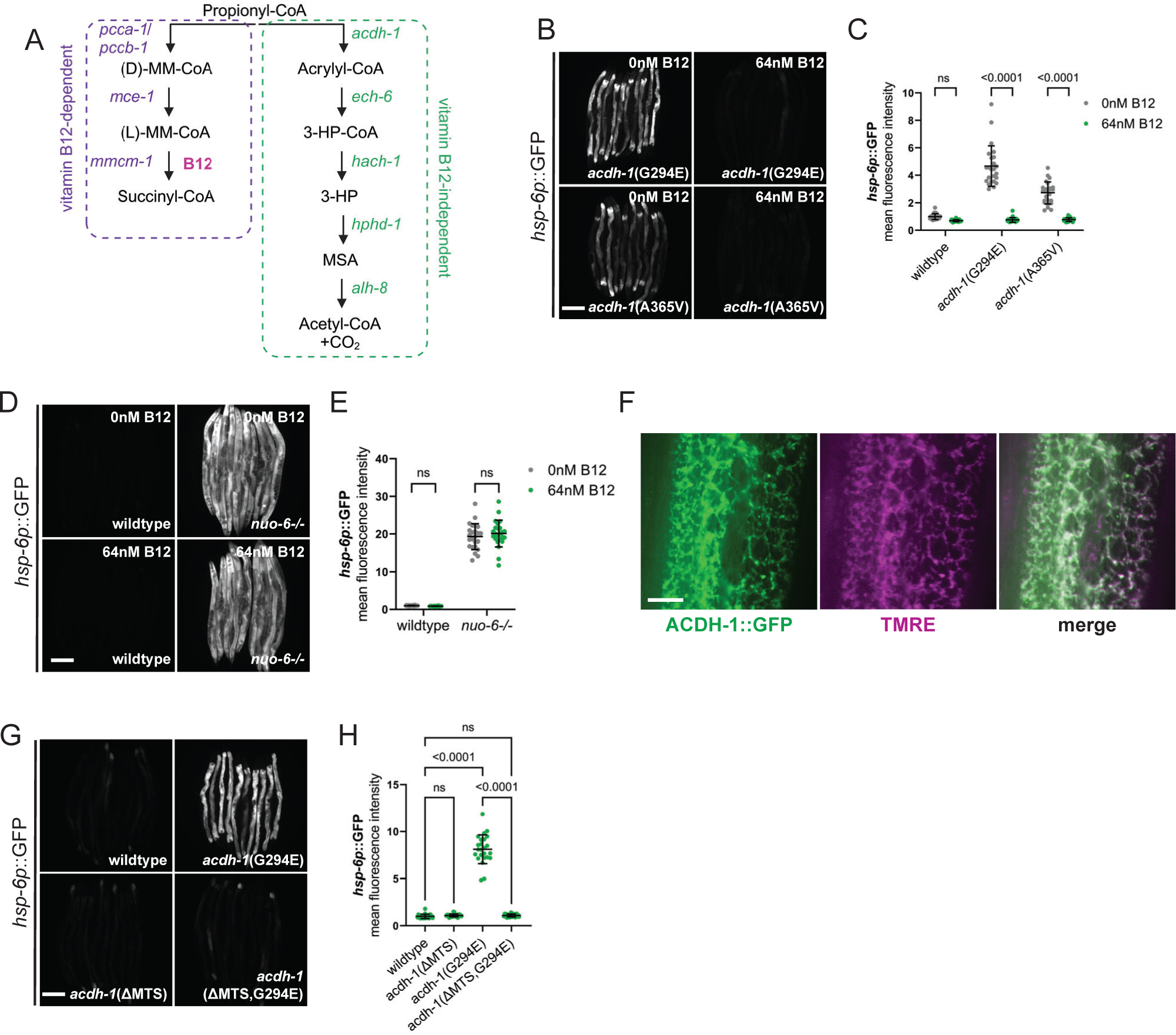
ACDH-1 is dependent upon vitamin B12 and required in the mitochondria to activate UPR^mt^. (A) Schematic of the parallel pathways known to metabolize the short chain propionic acid. (B) Fluorescence images of UPR^mt^ reporter (*hsp-6p*::GFP) activation in day 2 adult *acdh-1*(G294E) and *acdh-1*(A365V) animals supplemented with 0nM or 64nM vitamin B12. Scale bar 200 μm. (C) Relative mean fluorescence intensity quantification of *hsp-6p*::GFP in day 2 adult wildtype, *acdh-1*(G294E), and *acdh-1*(A365V) animals supplemented with 0nM or 64nM vitamin B12 (n=24, mean and SD shown, ordinary two-way ANOVA with Tukey’s multiple comparisons test). (D) Fluorescence images of UPR^mt^ reporter (*hsp-6p*::GFP) activation in day 2 adult wildtype and *nuo-6-/-* animals supplemented with 0nM or 64nM vitamin B12. Scale bar 200 μm. (E) Relative mean fluorescence intensity quantification of *hsp-6p*::GFP in day 2 adult wildtype and *nuo-6-/-* animals supplemented with 0nM or 64nM vitamin B12 (n=24, mean and SD shown, ordinary two-way ANOVA with Tukey’s multiple comparisons test). (F) Fluorescence images of the midsection of a wildtype animal expressing ACDH-1::GFP (green) stained with TMRE (magenta) to visualize mitochondria. GFP and TMRE co-localization shown in white in merged image. Scale bar 10 μm. (G) Fluorescence images of UPR^mt^ reporter (*hsp-6p*::GFP) activation in day 2 adult wildtype, *acdh-1*(G294E), *acdh-1*(ΔMTS), and *acdh-1*(ΔMTS,G294E) animals. Scale bar 200 μm. (H) Relative mean fluorescence intensity quantification of *hsp-6p*::GFP in day 2 adult wildtype, *acdh-1*(G294E), *acdh-1*(ΔMTS), and *acdh-1*(ΔMTS,G294E) animals (n=24, mean and SD shown, ordinary one-way ANOVA with Tukey’s multiple comparisons test).

### ACDH-1 is required in the mitochondria to activate UPR^mt^

Next, we aimed to identify where ACDH-1 functions in the cell to trigger UPR^mt^. ACDH-1 carries a mitochondrial targeting sequence and is predicted to localize to mitochondria ^35^. Confirming this, GFP-tagged ACDH-1 (allele mpt161, denoted as ACDH-1::GFP) we generated using CRISPR localizes to mitochondria as evidenced by colocalization with the mitochondrial specific dye, TMRE (Figure 4F). Removal of the mitochondrial targeting sequence in *acdh-1*(G294E) animals (*acdh-1*(ΔMTS,G294E) completely abolishes UPR^mt^ reporter activity (Figures 4G, 4H, and S4H). These data suggest that ACDH-1 is required in the mitochondria to mediate UPR^mt^.

### Acrylyl-CoA, an intermediate metabolite downstream of ACDH-1 and upstream of ECH-6, likely signals UPR^mt^ activation

The B12-independent propionate shunt pathway, initiated by ACDH-1, involves a series of enzymatic reactions that transform propionyl-CoA into acetyl-CoA (reference Figure 4A) ^29,36^. We aimed to identify the specific step in the propionate shunt pathway responsible for ACDH-1-dependent UPR^mt^. First, we revisited the strains from our forward genetic screen to determine whether we had recovered mutations in other enzymes within this pathway. We identified a clear loss-of-function mutation in *hach-1*, a gene encoding a hydroxyacyl-CoA hydrolase, that operates further downstream in the shunt pathway, in strain mpt138 (splice acceptor mutation chr III:5556870a>t). To validate the role of *hach-1* in UPR^mt^, we introduced the *hach-1* mutation into a clean genetic background using CRISPR (allele mpt152, denoted as *hach-1*-/-). This was sufficient to activate robust UPR^mt^ only in the intestine (Figures 5A, 5B, and S5A). This outcome suggests that the accumulation of a metabolite between ACDH-1 and HACH-1 triggers UPR^mt^. To further validate this hypothesis, we examined whether *hach-1*-/-induced UPR^mt^ is dependent on *acdh-1*. Indeed, UPR^mt^ reporter activation by *hach-1-/-* was completely abolished in the absence of *acdh-1* (Figures 5C, 5D, and S5B).

**Figure 5:**
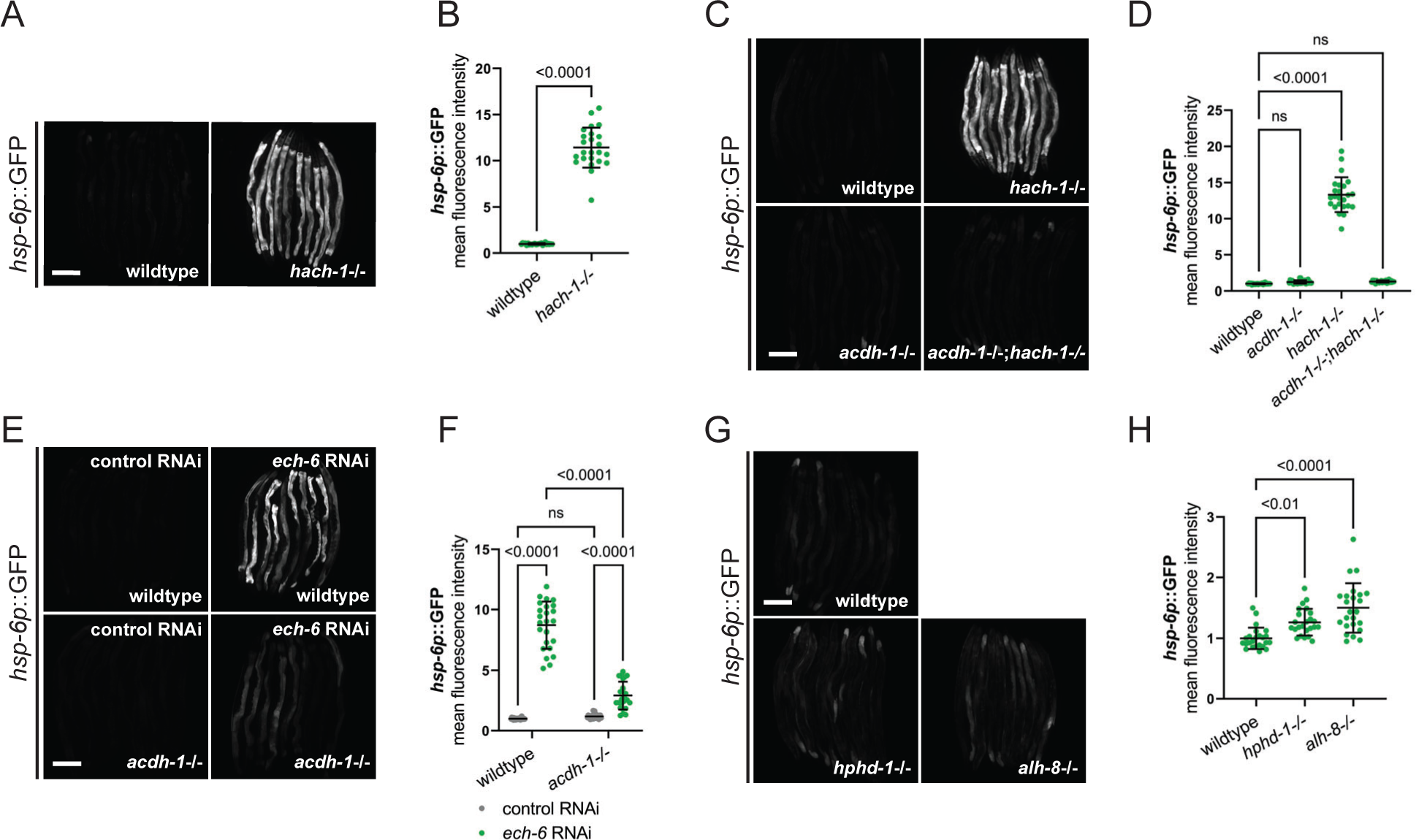
Acrylyl-CoA, an intermediate metabolite downstream of ACDH-1 and upstream of ECH-6, likely signals UPRmt activation. (A) Fluorescence images of UPR^mt^ reporter (*hsp-6p*::GFP) activation in day 2 adult wildtype and *hach-1* loss-of-function *(hach-1-/-*) animals. Scale bar 200 μm. (B) Relative mean fluorescence intensity quantification of *hsp-6p*::GFP in day 2 adult wildtype and *hach-1-/-* animals (n=24, mean and SD shown, unpaired t-test). (C) Fluorescence images of UPR^mt^ reporter (*hsp-6p*::GFP) activation in day 2 adult wildtype, *hach-1-/-*, *acdh-1-/-* and *acdh-1-/-;hach-1-/-* animals. Scale bar 200 μm. (D) Relative mean fluorescence intensity quantification of *hsp-6p*::GFP in day 2 adult wildtype *hach-1-/-*, *acdh-1-/-*, and *acdh-1-/-;hach-1-/-* double mutant animals (n=24, mean and SD shown, ordinary one-way ANOVA with Tukey’s multiple comparisons test). (E) Fluorescence images of UPR^mt^ reporter (*hsp-6p*::GFP) activation in day 2 adult wildtype and *acdh-1-/-* animals on control and *ech-6* RNAi. Scale bar 200 μm. (F) Relative mean fluorescence intensity quantification of *hsp-6p*::GFP in day 2 adult wildtype and *acdh-1-/-* animals on control and *ech-6* RNAi (n=24, mean and SD shown, ordinary two-way ANOVA with Tukey’s multiple comparisons test). (G) Fluorescence images of UPR^mt^ reporter (*hsp-6p*::GFP) activation in day 2 adult wildtype, *hphd-1* loss-of-function *(hphd-1-/-*), and *alh-8* loss-of-function (*alh-8-/-*) animals. Scale bar 200 μm. (H) Relative mean fluorescence intensity quantification of *hsp-6p*::GFP in day 2 adult wildtype, *hphd-1-/-*, and *alh-8-/-* animals (n=24, mean and SD shown, ordinary one-way ANOVA with Tukey’s multiple comparisons test).

Only one enzyme, an enoyl-CoA hydratase called ECH-6, is positioned between ACDH-1 and HACH-1 in this pathway ^29^. ECH-6 converts acrylyl-CoA, a highly reactive and unstable trans-enoyl-CoA, into 3-hydroxypropionyl-CoA. To pinpoint the precise metabolic step triggering ACDH-1 dependent UPR^mt^, we attempted to knockout *ech-6* using CRISPR. However, we could not recover homozygous loss-of-function mutants, suggesting that *ech-6* is essential. Consequently, we employed RNAi to knockdown *ech-6*, which resulted in robust UPR^mt^ activation in an *acdh-1* dependent manner (Figures 5E, 5F, and S5C). Notably, loss-of-function mutations in the downstream components of the propionate shunt pathway, *hphd-1* and *alh-8* (alleles: ok3580 and ww48 respectively, denoted as *hphd-1*-/-and *alh-8*-/-), had no impact on the UPR^mt^ reporter (Figure 5G, 5H, and S5D). Collectively, these findings suggest that the accumulation of a short-chain fatty acid intermediate metabolite, acrylyl-CoA, positioned between ACDH-1 and ECH-6, serves as a signal for UPR^mt^ activation.

### HOE-1 functions downstream of ACDH-1

We have established that both HOE-1 and ACDH-1 activate post-developmental, intestinal-specific UPR^mt^. Therefore, our next objective was to investigate whether they function within the same genetic pathway. Our findings reveal that the complete loss of *acdh-1* or the targeted loss of *acdh-1* within mitochondria strongly attenuates UPR^mt^ in *glp-1*(ts);*hoe-1*(OE) animals (Figures 6A–D, S6A, and S6B). Moreover, akin to the ACDH-1-dependent UPR^mt^, the UPR^mt^ triggered by HOE-1 overexpression in animals lacking a germline is also inhibited by vitamin B12 supplementation (Figures 6E, 6F, and S6C).

**Figure 6:**
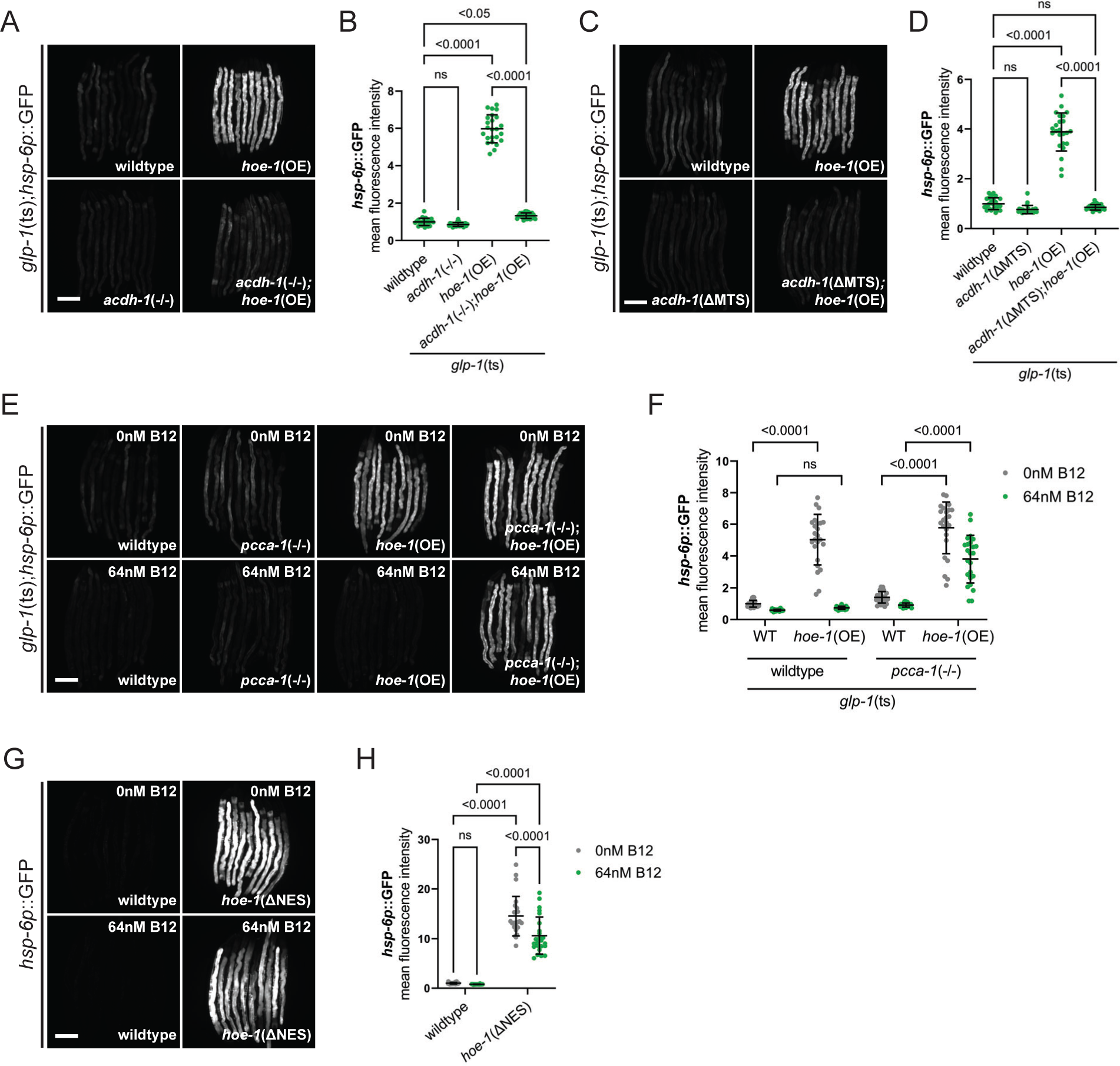
HOE-1 functions downstream of ACDH-1. (A) Fluorescence images of UPR^mt^ reporter (*hsp-6p*::GFP) activation in day 2 adult wildtype, *hoe-1*(OE), *acdh-1*-/-, and *acdh-1*-/-;*hoe-1*(OE) animals all of which are in a *glp-1*(ts) background at the restrictive temperature (25°C). Scale bar 200 μm. (B) Relative mean fluorescence intensity quantification of *hsp-6p*::GFP in day 2 adult wildtype, *hoe-1*(OE), *acdh-1*-/-, and *acdh-1*-/-;*hoe-1*(OE) animals all of which are in a *glp-1*(ts) background at the restrictive temperature (25°C) (n=24, mean and SD shown, ordinary one-way ANOVA with Tukey’s multiple comparisons test). (C) Fluorescence images of UPR^mt^ reporter (*hsp-6p*::GFP) activation in day 2 adult wildtype, *hoe-1*(OE), *acdh-1*(ΔMTS), and *acdh-1*(ΔMTS);*hoe-1*(OE) animals all of which are in a *glp-1*(ts) background at the restrictive temperature. Scale bar 200 μm. (D) Relative mean fluorescence intensity quantification of *hsp-6p*::GFP in day 2 adult wildtype, *hoe-1*(OE), *acdh-1*(ΔMTS), and *acdh-1*(ΔMTS);*hoe-1*(OE) animals all of which are in a *glp-1*(ts) background at the restrictive temperature (25°C) (n=24, mean and SD shown, ordinary one-way ANOVA with Tukey’s multiple comparisons test). (E) Fluorescence images of UPR^mt^ reporter (*hsp-6p*::GFP) activation in day 2 adult wildtype, *pcca-1* loss-of-function mutant (*pcca-1*-/-), *hoe-1*(OE), and *pcca-1*-/-;*hoe-1*(OE) animals all of which are in in a *glp-1*(ts) background at the restrictive temperature (25°C), supplemented with either 0 or 64 nM vitamin B12. Scale bar 200 μm. (F) Relative mean fluorescence intensity quantification of *hsp-6p*::GFP in day 2 adult wildtype, *pcca-1*-/-, *hoe-1*(OE), and *pcca-1*-/-;*hoe-1*(OE) animals all of which are in in a *glp-1*(ts) background at the restrictive temperature (25°C), supplemented with either 0 or 64 nM vitamin B12 (n=24, mean and SD shown, ordinary two-way ANOVA with Tukey’s multiple comparisons test). (G) Fluorescence images of UPR^mt^ reporter (*hsp-6p*::GFP) activation in day 2 adult wildtype and *hoe-1*(ΔNES) animals supplemented with 0nM and 64nM vitamin B12. Scale bar 200 μm. (H) Relative mean fluorescence intensity quantification of *hsp-6p*::GFP in day 2 adult wildtype and *hoe-1*(ΔNES) animals supplemented with 0nM and 64nM vitamin B12 (n=24, mean and SD shown, ordinary two-way ANOVA with Tukey’s multiple comparisons test).

In *C. elegans*, vitamin B12 is required as a cofactor for two enzymes: methylmalonyl-CoA mutase of the vitamin B12-dependent propionate metabolism pathway (parallel to the propionate shunt pathway), and methionine synthase of the one carbon cycle ^36^. Thus, while vitamin B12 is sufficient to attenuate *acdh-1* expression, it is also possible that vitamin B12 supplementation may indirectly affect UPR^mt^ via rescue of the one carbon cycle. To validate that this suppression by vitamin B12 supplementation is mediated through *acdh-1* directly, we repeated these experiments in animals lacking PCCA-1, the alpha subunit of the propionyl coenzyme A carboxylase that functions in the vitamin B12-dependent propionate metabolism pathway ^37^. In the absence of PCCA-1, *acdh-1* expression persists even in the presence of elevated vitamin B12 levels ^33^. Our findings indicate that the loss of *pcca-1* restores UPR^mt^ activation in *glp-1*(ts);*hoe-1*(OE) animals, when vitamin B12 is supplemented (Figures 6E, 6F, and S6C). Collectively, the ACDH-1 and vitamin B12 dependency of HOE-1-induced UPR^mt^ establishes ACDH-1 and HOE-1 as components of the same genetic pathway.

Next, we aimed to elucidate the epistatic relationship between ACDH-1 and HOE-1 by investigating their interdependence. Knockdown of *hoe-1* inhibited *acdh-1* expression, precluding us from assessing whether ACDH-1-driven UPR^mt^ relies on *hoe-1* (Figures S6D and S6E). However, we found that UPR^mt^ in *hoe-1*(ΔNES) animals, is still robustly activated despite vitamin B12 supplementation (Figure 6G, 6H, and S6F). These findings position HOE-1 downstream of ACDH-1.

### Germline non-autonomously regulates ACDH-1 pathway activity

A model emerges from our data for UPR^mt^ induction wherein a non-functional germline cell non-autonomously influences an ACDH-1-dependent pathway to enhance HOE-1 activity. It is still unclear, however, through what mechanism the germline impacts the ACDH-1 pathway. To address this, we examined the *acdh-1* transcriptional reporter in *glp-1*(ts) animals, which revealed increased *acdh-*1 in *glp-1*(ts) animals at an early adult stage (Figure 7A, 7B, and S7A). Considering that reducing *ech-6* activity produces the same outcome as activating *acdh-1*, we also assessed ECH-6 protein levels using a GFP-tagged CRISPR strain (allele mpt188, denoted as ECH-6::GFP). We observed a substantial decrease in ECH-6 protein levels in *glp-1(ts)* animals (Figure 7C, 7D and S7B). Together, these data show that the germline non-autonomously alters the propionate shunt pathway by upregulating *acdh-1* expression and reducing ECH-6 levels.

**Figure 7:**
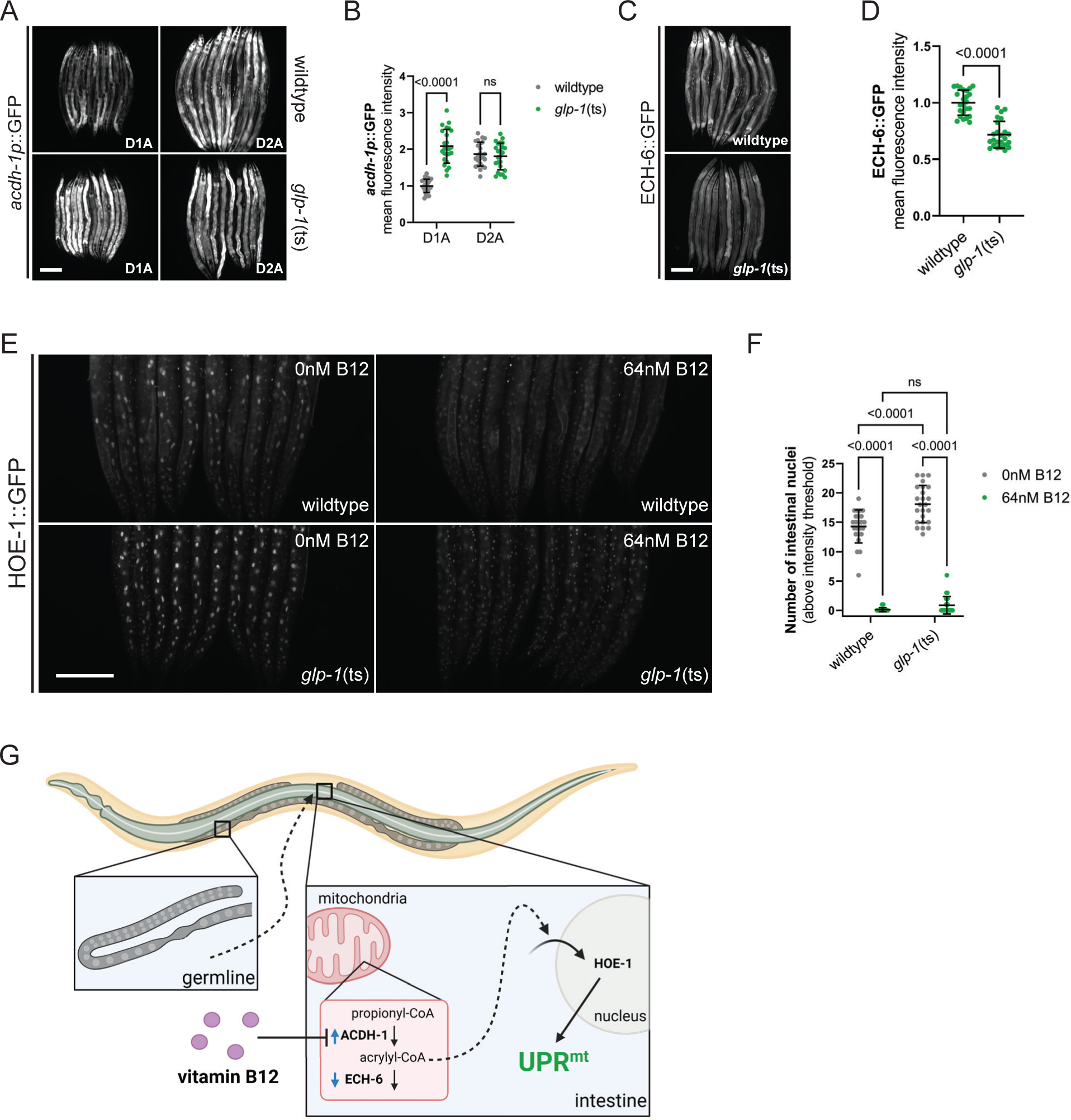
The germline non-cell autonomously regulates ACDH-1 pathway activity, influencing nuclear accumulation of HOE-1. (A) Fluorescence images of *acdh-1* transcriptional reporter (*acdh-1p*::GFP) activation in day 1 (D1) and day 2 (D2) adult wildtype and *glp-1*(ts) animals at the restrictive temperature (25°C). Scale bar 200 μm. (B) Relative mean fluorescence intensity quantification of *acdh-1p*::GFP activation in day 1 (D1) and day 2 (D2) adult wildtype and *glp-1*(ts) animals at the restrictive temperature (25°C) (n=24, mean and SD shown, ordinary two-way ANOVA with Tukey’s multiple comparisons test). (C) Fluorescence images of ECH-6 protein reporter (ECH-6::GFP) activation in day 2 adult wildtype and *glp-1*(ts) animals at the restrictive temperature (25°C). Scale bar 200 μm. (D) Relative mean fluorescence intensity quantification of ECH-6::GFP activation in day 2 adult wildtype and *glp-1*(ts) animals at the restrictive temperature (25°C) (n=24, mean and SD shown, ordinary one-way ANOVA with Tukey’s multiple comparisons test). (E) Fluorescence images of HOE-1 protein reporter (HOE-1::GFP) in wildtype and *glp-1*(ts) animals at the restrictive temperature (25°C) supplemented with 0 or 64 nM vitamin B12 (zoomed in on posterior half of animals). Scale bar 200 μm. (F) Number of intestinal nuclei per animal with a HOE-1::GFP fluorescence intensity exceeding a pixel intensity threshold of 100 in wildtype and *glp-1*(ts) animals at the restrictive temperature (25°C) supplemented with 0 or 64 nM vitamin B12 (n=24, mean and SD shown, ordinary two-way ANOVA with Tukey’s multiple comparisons test). (G) Compromised germline integrity cell non-autonomously regulates the mitochondrial propionate shunt pathway resulting in accumulation of acrylyl-CoA which in turn drives increased nuclear accumulation of HOE-1 triggering intestinal specific UPR^mt^ (figure generated using BioRender).

### Nonfunctional germline promotes nuclear accumulation of HOE-1

The second question raised by our model pertains to how the germline influences HOE-1 activity. Given that the loss of germline induced by *glp-1*(ts) robustly activates UPR^mt^ in combination with *hoe-1*(OE), one possibility is that *glp-1*(ts) facilitates nuclear accumulation of HOE-1, elevating HOE-1 levels in the nucleus beyond a critical threshold required to trigger UPR^mt^. In line with this possibility, *glp-1*(ts) animals exhibit higher nuclear localization of HOE-1::GFP compared to wildtype animals (Figure 7E, 7F, and S7C). Notably, supplementation with vitamin B12 drastically reduces HOE-1 nuclear levels in both wildtype and *glp-1*(ts) animals (Figure 7E, 7F, and S7C). Taken together, these data support our model, indicating that the germline modulates nuclear levels of HOE-1 through ACDH-1 (Figure 7G).

## DISCUSSION

The capacity of somatic cells to sense, integrate, and respond to reproductive cues forms the basis of metabolic adaptation in females. Here we show how reproductive status and dietary availability of vitamin B12 can work together to control UPR^mt^, a mitochondrial quality control pathway. Mechanistically, ACDH-1, a mitochondrially localized acyl-CoA dehydrogenase, serves as the critical link between germline status and HOE-1 regulation in the soma. Specifically, the state of the germline regulates the ACDH-1-dependent propionate shunt pathway in the soma. This pathway, via a metabolic intermediate, regulates the movement of HOE-1 into the nucleus, leading to the activation of UPR^mt^. Our findings uncover fundamental principles by which cells synthesize physiological signals to regulate mitochondrial quality control.

ELAC2, the mammalian counterpart of HOE-1, functions as an RNA endonuclease primarily associated with an essential housekeeping task in tRNA processing ^12–15^. Therefore, HOE-1’s unexpected role in selectively regulating mitochondrial function and UPR^mt^ is both surprising and potentially insightful. Mutations in ELAC2 are linked to prostate cancer and are known to cause hypertrophic cardiomyopathy ^15–22^. The cellular logic underlying ELAC2’s involvement in these diseases has remained elusive. While one hypothesis posits that these conditions stem from mitochondrial dysfunction due to compromised tRNA processing in mitochondria, our findings suggest an alternate paradigm. Specifically, mutations in ELAC2 may induce mitochondrial dysfunction, in part, by disrupting the nuclear-localized function of HOE-1 in the regulation of mitochondrial quality control. Investigating this paradigm in the future, particularly within mammalian systems, holds the potential to yield valuable insights into ELAC2’s disease mechanisms.

Interestingly, although ELAC2 has a well-established role in processing precursor tRNAs, emerging evidence indicates that ELAC2 can cleave mature tRNAs, generating tRNA fragments with biological activity ^38,39^. Additionally, ELAC2 exhibits non-tRNA target cleavage, exemplified by its processing of MALAT1, a long noncoding RNA implicated in many cancers ^40^. Collectively, these findings suggest ELAC2 has an essential role in the fundamental function of tRNA maturation and hint at additional roles in RNA biology. It will be interesting to determine which HOE-1 RNA substrates induce UPR^mt^ and to delineate their mechanism of action.

In the broader context of exploring HOE-1’s role in regulating mitochondrial quality control, it is crucial to understand the physiologically relevant conditions governing HOE-1 activity. Within this framework, our findings reveal the critical involvement of the germline’s functional status as a key cell non-autonomous factor influencing the dynamics of HOE-1’s nuclear localization. Mechanistically, the germline achieves this by activating the ACDH-1-dependent propionate shunt pathway localized in the mitochondria within somatic cells. ACDH-1, in turn, generates acrylyl-CoA, which our genetic analysis suggests serves as a signaling factor promoting the nuclear localization of HOE-1. Interestingly, because the expression of *acdh-1* is influenced by the availability of vitamin B12 ^29,33^, the germline’s ability to modulate HOE-1-induced UPR^mt^ is dictated by the levels of this essential dietary micronutrient. Thus, the ACDH-1 pathway integrates information about both the germline status and vitamin B12 availability to regulate HOE-1-mediated mitochondrial quality control.

Acrylyl-CoA is an unstable and highly reactive metabolic intermediate in the propionate shunt pathway ^41^. In metazoans, it has predominantly been studied in the context of disease arising from loss-of-function mutations in ECHS1, the mammalian homolog of ECH-6 ^42–47^. Interestingly, these diseases exhibit phenotypic similarities to Leigh syndrome, typically associated with mitochondrial dysfunction ^42,47^. While detecting acrylyl-CoA is challenging, diseases linked to EHCS1 deficiency can be diagnosed by evaluating acrylyl-CoA conjugate levels in patients. It is hypothesized that acrylyl-CoA toxicity is the basis for pathogenicity in ECHS1-defective patients ^42,47^. Our study provides evidence for acrylyl-CoA’s role as a signaling molecule rather than a mere toxin. In this sense, acrylyl-CoA is akin to reactive oxygen species (ROS), acting as signaling molecules when produced in a controlled manner but turning toxic at pathologically high levels ^48^. The concept of acrylyl-CoA playing a signaling role is not unprecedented as it is part of a larger family of acyl-CoAs such as acetyl-CoA and crotonyl-CoA that have well-established and emerging functions as signaling agents ^49,50^. Exploring the signaling role of acrylyl-CoA by identifying its molecular targets promises to be fruitful avenue for further investigation.

The impact of germline status on the soma has been most extensively studied within the context of aging in *C. elegans* ^51^. The absence of a germline is recognized for its remarkable ability to significantly prolong lifespan ^52,53^. The impact on lifespan seems to be mediated, at least partly, by the regulation of somatic stress responses ^54–56^. Reproductive maturity coincides with a significant impairment in the ability to initiate various stress responses ^57^. However, adult animals lacking a functional germline maintain the ability to robustly activate the heat shock response ^57^. Here, we discovered that instead of merely preserving the capacity of adults to induce UPR^mt^, the absence of a functional germline actively facilitates UPR^mt^ induction. This unique and unusual finding suggests potential differences in the mechanisms involved. The identification and subsequent comparison of these mechanisms promises to provide insights into the intricate interplay between germline status and somatic stress responses.

Although the germline plays a significant role in regulating HOE-1 dependent UPR^mt^, we discovered that it is highly sensitive to vitamin B12 levels. Why has the regulation of UPR^mt^ evolved to be responsive to vitamin B12? One possible explanation is that vitamin B12 is crucial for mitochondrial function, especially in the context of reproduction. Consistent with this notion, several studies have reported the impact of vitamin B12 on mitochondria, with one recent finding demonstrating that vitamin B12 affects reproductive aging through its effect on mitochondrial positioning in oocytes ^58–62^. Moreover, the importance of vitamin B12 in human reproduction has long been recognized while the molecular mechanisms are still being uncovered. It will be insightful to further investigate the specific connection between vitamin B12 and mitochondria.

## ACKNOWLEDGEMENTS

We thank WormBase for invaluable tools and information used to plan and execute the research described. Some strains were provided by the CGC, which is funded by NIH Office of Research Infrastructure Programs (P40 OD010440). RNA and whole genome sequencing were conducted by the Vanderbilt University Medical Center’s VANTAGE Core (NIH 1U24OD035523-01). Some schematics were generated using BioRender. This work was generously supported by R01 GM123260 (MRP), R35 GM145378 (MRP), pilot grant from the Evolutionary Studies at Vanderbilt (MRP). JPH, LKG, and SHS are supported by the Training Program in Environmental Toxicology (T32ES007028). LKG is supported by National Defense Science & Engineering Graduate Fellowship through the US Department of Defense. SHS is supported by the Graduate Research Fellowship Program through the National Science Foundation.

## AUTHOR CONTRIBUTIONS

Conceptualization, J.P.H and M.R.P.; Methodology, J.P.H and M.R.P.; Validation, J.P.H., A.M.S., and H.R.; Formal Analysis, J.P.H. and L.K.G.; Investigation, J.P.H., N.H.D, A.M.S, L.K.G., H.R., and M.R.P.; Resources, J.P.H., H.R., S.H.S., and M.R.P.; Data Curation, J.P.H., and L.K.G.; Writing – Original Draft, J.P.H. and M.R.P.; Writing – Review & Editing, J.P.H., N.H.D, A.M.S, L.K.G., H.R., S.H.S., and M.R.P.; Visualization, J.P.H. and L.K.G.; Supervision, J.P.H. and M.R.P.; Project Administration, J.P.H. and M.R.P.; Funding Acquisition, M.R.P.

## STAR METHODS

### RESOURCE AVAILABILITY

#### Lead Contact

Further information and requests for resources and reagents should be directed to the lead contact, Maulik R. Patel.

#### Materials Availability

Any reagent generated for this manuscript is available from the lead contact upon request.

#### Data and Code Availability

Processed data is provided as supporting documentation with this manuscript. All raw and processed data and analysis codes will be made publicly available upon final publication. Any additional information required to reanalyze the data reported in this paper is available from the lead contact upon request.

## METHOD DETAILS

### *C. elegans* Maintenance

Worms were grown on nematode growth media (NGM) seeded with OP50 *E. coli* bacteria and maintained at 20°C. Strains containing temperature sensitive mutants were maintained at 16°C. All experiments were conducted under these conditions unless otherwise stated.

### Bacterial Strains

Bacterial strain OP50 *E. coli* was obtained from the *Caenorhabditis* Genetics Center (RRID: WBStrain00041969). RNAi bacterial strains are all in the *E. coli* strain HT115 as part of the Ahringer RNAi Library [S1].

### Mutant and Transgenic Lines

A full list of mutant and transgenic worm strains used can be found in the Key Resource Table. All new mutant and transgenic strains generated via CRISPR/Cas9 for this study were confirmed by Sanger sequencing.

### CRISPR/Cas9

CRISPR was conducted as previously described using Alt-R S.p. Cas9 Nuclease V3 (IDT #1081058) and tracrRNA (IDT #1072532) [S2]. However, instead of using the *rol-6* plasmid, we used *dpy-10* endogenous editing as a co-injection marker as previously described [S3]. Once the edit of interest was recovered, the *dpy-10* co-marker was outcrossed using a wildtype (N2, RRID: WBStrain00000001) background. A complete list of crRNA and repair template sequences purchased from IDT can be found in Table S1.

### Genetic Crosses

Strains resulting from genetic crosses were generated by crossing 15-20 heterozygous males of a given strain to 5-8 larval stage 4 (L4) hermaphrodites of another strain (heterozygous males were first generated by crossing wildtype N2 males to L4 hermaphrodites of a strain). F1 generation L4s were subsequently cloned out from cross plates. Once F2 progeny were laid, F1s were genotyped/screened for allele(s) of interest. F2 progeny were cloned out from F1s heterozygotes, and once F3 progeny were laid, F2s were genotyped/screened for homozygosity of alleles of interest. All genotyping primers were purchased from IDT and can be found in Table S1.

### RNA Extraction for RNA Sequencing

Stage synchronized animals were grown from embryo at 25°C for 72 hours. 800 adult animals were collected for each replicate and transferred into 1 ml of M9 Buffer in a 1.5 mL microcentrifuge tube. Worms were pelleted at 500xg for 1 minute, supernatant removed and then washed once with 1 ml of fresh M9, and then once with 1 ml of M9 supplemented with 0.01% Tween-20 following the same spin and supernatant removal. After removing as much of the last supernatant as possible, 250 ul of Qiazol was added to each sample and snap-frozen in liquid nitrogen. Samples were then run through 3 freeze thaw cycles: 37°C bead bath for 2 min followed by liquid nitrogen for 1 minute. Following freeze thaw cycles, samples were completely thawed and incubated at room temperature for 5 minutes. 50 ul of chloroform was added to each sample, vigorously vortexed for 30 s and incubated at room temperature for 3 minutes. Then, samples were centrifuged at 4°C for 15 minutes at 12,000xg for 15 min. Following the spin, 125 ul of the upper aqueous phase was transferred to a fresh 1.5 ml tube. 187.5 ul of 100% molecular grade ethanol was then added to each sample and pipetted up and down to mix. Samples were then transferred to Qiagen RNeasy Mini Kit columns, centrifuged at 8,000xg for 30 seconds, and flow through discarded. Then, 700 ul of RW1 buffer was added to column, centrifuged at 8,000xg for 30 seconds, and flow through discarded. 500 ul RPE buffer added to column, centrifuged at 8,000xg for 30 seconds, and flow through discarded. Then again 500 ul of RPE buffer was added, centrifuged at 12,000xg for 2 min, and flow through discarded. Columns were then transferred to fresh collection tubes, centrifuged at 13,000xg for 1 minute with tube lids open to dry columns. Samples were then eluted from column with 20 ul of RNase free water into a fresh 1.5 ml microcentrifuge tube by centrifuging at 13,000xg for 1 min.

Following RNA extraction, RNA samples were DNase treated with Turbo DNA-free Kit (Invitrogen #AM1907) following manufacturer’s directions. Briefly, 2 ul of 10x Turbo DNase Buffer and 1 ul of Turbo DNase were added to 20 ul RNA samples and gently mixed by pipetting up and down. Samples were then incubated in a 37°C bead bath for 20 min. Following incubation, 3 ul of resuspended Inactivation Reagent was added to each sample and mixed well by pipetting up and down. Samples were incubated at 25°C for 5 min. Then, samples were centrifuged at 10,000xg for 90 sec. The RNA containing supernatants were transferred to fresh tubes and stored at -80°C until RNA sequencing.

### RNA Sequencing

RNA quality control, library preparation, and RNA sequencing were conducted by the Vanderbilt University Medical Center Vanderbilt Technologies for Advanced Genomics (VANTAGE) core. RNA sample quality and integrity was assessed by fluorometry Qubit or Picogreen (for concentration) and by BioAnalyzer or TapeStation (for integrity), respectively. Stranded mRNA library preparation was conducted using NEBNext® Poly(A) selection. Sequencing was performed at Paired-End 150 bp on the Illumina NovaSeq 6000 targeting an average of 50M reads per sample.

### RNA Sequencing Analysis

An average of ∼40,000,000 reads were generated per sample and the Q30+ percentage was ∼92% on average. Initial read quality was assessed with FastQC. Paired end reads were trimmed using Trimmomatic-0.39 [S4]. The universal Illumina adapter was removed, and reads were trimmed with a 4:15 sliding window. STAR 2.5.4 was used to align reads to the genome and transcriptome (WBcel.235) using default parameters [S5]. Salmon was used to generate transcript pseudo counts with the following options: --validateMappings --recoverOrphans -- numBootstraps=30 --useVBOpt --seqBias --gcBias –writeUnmappedNames [S6]. The *C. elegans* Ensembl database was used to collate gene IDs. Tximport was used to import data from the salmon files, including read abundance, length, counts, counts from abundance [S7]. DESeq2 was used to generate a contrast and determine significantly different transcript levels [S8]. DESeq2 was also used to normalize counts to account for variation in library size.

### Gene Ontology Analysis

Gene Ontology Analysis was conducted using the STRING database [S9]. For Figure 1B, genes with a log_2_ fold change >1 and an adjusted p-value <0.05 were compiled for entry into the ‘multiple proteins’ feature of STRING.

### Fluorescence Microscopy

All whole animal imaging was done using a Zeiss Axio Zoom V16 stereo zoom microscope. For all imaging, worms were immobilized on 2% agar pads on microscope slides in ∼1 μl of 100 mM levamisole (ThermoFisher #AC187870100) and then coverslip applied.

### Fluorescence Image Analysis

For whole animal fluorescence intensity quantification, mean fluorescence intensity of each individual animal was determined by tracing the outline of each worm and then using the ‘measure’ feature in ImageJ/Fiji (which calculates the average pixel intensity by dividing the sum total of fluorescence intensity by the total number of pixels within bounds of the trace). For HOE-1::GFP image analysis (Figure 7E, 7F), the ‘threshold’ feature of ImageJ/Fiji was used to count the number of gut cell nuclei that were saturated at a pixel intensity threshold of 100.

### EMS Mutagenesis Screen

Mutagenesis was conducted as previously described [S10]. Briefly, zcIs13 animals were washed off of four, 60 mm plates (containing at least a few hundred larval stage 4 (L4) animals total) with sterile M9 buffer. Worms were collected in a 15 ml conical tube which was centrifuged at 1500xg for 3 min to pellet worms. Supernatant was removed by aspiration. The worm pellet was resuspended in 2 ml of M9. In a separate tube, 20 μl of ethyl methanesulfonate (EMS) was added to 2 ml of M9 buffer then vortexed. The 2 ml of EMS supplemented M9 buffer was added to the 2 ml worm suspension (final concentration of EMS in solution is ∼50 mM). The worms were incubated for 4 hours at room temperature constantly mixing on a nutator. After incubation, worms were pelleted by centrifugation at 1500xg for 3 minutes, supernatant was removed. Worms were then washed 3 times with 5 ml of fresh M9 buffer by sequential centrifugation at 1500xg for 3 minutes, removal of supernatant, and addition of M9. After the final wash, worms were resuspended in a minimal volume of M9 buffer (∼200 – 500 μl) and then transferred to NGM plates seeded with OP50 bacteria. Worms were allowed to recover for 2 hours, then 5 L4 animals (considered P0 generation) were transferred to separate OP50 seeded plates (10 – 20 plates of 5 L4s each per round of mutagenesis). 3 – 5 days following mutagenesis, L4 or young adult F1 animals were cloned out to individual 35 mm plates (500 animals per day over a three-day period; ∼1500 animals per round of mutagenesis). A minimum of 6000 F1 animals were cloned in total. Once F1 containing plates had a majority of progeny reach adulthood (3 – 5 days post F1 cloning), F2 populations were screened for UPR^mt^ activation. Special attention was paid to plates that had UPR^mt^ activation in adult animals but not larval animals as well as UPR^mt^ activation only in the intestine. Putative ‘hits’ (UPR^mt^ intestinal activation and in adults only) were transferred to individual plates. These populations were subsequently assessed to confirm that UPR^mt^ was only activated in adults and that the causal mutation was homozygous (all offspring exhibited UPR^mt^ activation). These were the populations that were kept as true hits (which were mpt134, mpt135, mpt136, mpt137, mpt138, and mpt140). Six rounds of backcrossing were conducted for each mutant to remove the majority of passenger mutations caused by EMS.

### DNA Extraction for Whole Genome Sequencing

DNA extraction was conducted as previously described [S11, S12]. *C. elegans* of each mutant recovered from the screen were collected from a densely populated 150 mm plates by washing the animals off the plate with M9 buffer and collected in 15 ml conical tubes. Worms were pelleted by centrifugation at 450 x g for 3 min. The supernatant was discarded, worms were washed three times with 15 mL of M9 buffer and pelleted at 450 x g. Worms were then washed once with 15 mL of milliQ water, spun down, and the supernatant was discarded. Total worm pellet collected was ∼1 ml. Animals were resuspended in 2 mL of worm lysis buffer (0.1M Tris-Cl pH 8.5, 0.1M NaCl, 50 mM EDTA pH 8.0, 1% SDS) and supplemented with 100 µL of Proteinase K 20 mg/mL (ThermoFisher, 25530049). These were mixed by inversion and incubated for 1 hour at 62°C. After incubation, samples were supplemented with 400 µL 5 M NaCl and mixed by inversion. Samples were further supplemented with 400 µL of CTAB solution (10% CTAB, 4% NaCl) and incubated for 10 min at 37°C. To that mixture, 2 ml of chloroform (Sigma Aldrich, C2432) was added and vortexed vigorously for ten seconds. Phase separation was achieved by centrifuging at 2000 x g for 10 min at room temperature. The aqueous phase was recovered and mixed with 2 mL of Phenol:Chloroform:Isoamyl Alcohol (25:24:1) saturated with 10 mM Tris, pH 8.0, 1 mM EDTA (Sigma Aldrich, P3803). This was vortexed vigorously for 10 seconds and phase separation was achieved by centrifugation at 2000 x g for 10 min at room temperature. The aqueous phase was transferred to a 2 ml microcentrifuge tube, supplemented with 0.6 volumes of -20°C isopropanol, and mixed by inversion. DNA was precipitated by chilling samples at -20°C for 30 min followed by centrifugation at 13,000 x g for 5 min at room temperature. The pellet was washed twice with ice cold 70% ethanol and then resuspended in 200 µL TE. The 200 µL DNA sample was supplemented with 20 µL of RNase A (Thermo Fisher, EN053) and mixed by flicking the tube and inverting several times. The samples were incubated for 2 hours at 37°C. The sample was then supplemented with 20 µL of 20% SDS, 10 µL of 0.5 M EDTA pH 8.0, and 20 µL of Proteinase K. The samples were incubated for 1 hour at 62°C. Following incubation, samples were supplemented with 40 µL 10 M ammonium acetate and mixed. The DNA was extracted with an equal volume of Phenol:Chloroform:Isoamyl Alcohol saturated with TE pH 8.0. The sample was mixed vigorously by vortexing and phase separation was achieved by centrifuging 5 min at 5000 x g. The aqueous phase was recovered and was extracted again with an equal volume of chloroform. The sample was mixed vigorously by vortexing and phase separation was achieved by centrifuging 5 min at 5000 x g. DNA was precipitated by adding 2 volumes of -20°C 100% ethanol. The sample was mixed by inversion several times and allowed to chill at -20°C for 1 hour. The DNA was pelleted by centrifugation at 13,000 x g for 5 min at room temperature. The DNA pellet was washed twice with ice cold 70% ethanol and resuspended in 50 µL nuclease free water.

### Whole Genome Sequencing

DNA quality control, library preparation, and whole genome sequencing were conducted by the Vanderbilt University Medical Center Vanderbilt Technologies for Advanced Genomics (VANTAGE) core. DNA quality and content was assessed via Qubit. Then library preparation was conducted (Illumina Nextera DNA Flex). WGS was conducted on Illumina NovaSeq6000 with PE150 targeting 30x coverage per sample. Initial analysis to generate variant call files was conducted in Illumina’s BaseSpace program.

### Whole Genome Analysis and Variant Identification

We filtered the variants first by eliminating variants with an allele count >4 and then those present in <85% of the reads. This left us with high confidence variants as variants that are causal mutations should be present in nearly all reads and due to the outcrossing strategy. Remaining variants were manually identified and annotated in IGV to identify candidate causal mutations.

### TMRE Staining

A 10 mM stock solution of TMRE (Sigma Allrich #87917) in DMSO was first prepared. 100 μl of 1 μM TMRE solution (diluted from stock in sterile water) was added to the top of a 1-day seeded OP50 NGM plate and allowed to dry in the dark for 2 hours. Then, larval stage 4 (L4) worms were transferred to TMRE plates and incubated at 20°C in the dark for 24 hours. Following incubation, worms were transferred to a non-TMRE containing OP50 seeded NGM plate and allowed to recover from the dye (i.e. remove any excess dye from intestinal lumen, body wall, etc.) for 1 hour in the dark. Following recovery, animals were imaged immediately.

### Vitamin B12 Supplementation

During NGM plate preparation, a stock of 1 mM vitamin B12 in water was supplemented to molten agar (once cooled to <55°C) to a final concentration of 64 nM.

### RNAi

RNAi by feeding was conducted as previously described [S13]. Briefly, RNAi clones were grown overnight from single colony in 2 ml liquid culture of LB supplemented with 50 μg/ml ampicillin. To make 16 RNAi plates, 50 ml of LB supplemented with 50 μg/ml ampicillin and inoculated with 500 μl of overnight culture, then incubated while shaking at 37 °C for 4–5 hours (to an OD550-600 of ∼0.8). Then, to induce expression of double stranded RNA, cultures were supplemented with an additional 50 ml of LB supplemented with 50 μg/ml ampicillin and 4 mM IPTG and then continued incubating while shaking at 37 °C for 4 hours. Following incubation, bacteria were pelleted by centrifugation at 3900 rpm for 6 min. Supernatant was decanted and pellets were gently resuspended in 4 ml of LB supplemented with 8 mM IPTG. 250 μl of resuspension was seeded onto standard NGM plates containing 1 mM IPTG. The bacterial background used for RNAi, *E. coli* HT115, has higher B12 levels than OP50. Thus, NGM plates were prepared with soy peptone to reduce accessible vitamin B12. Plates were left to dry overnight and then used within 1 week. Bacterial RNAi feeder strains were from Ahringer RNAi Feeding Library, grown from single colony and identity confirmed by Sanger sequencing. Ech-6 (T05G5.6) hoe-1 (E04A4.4).

### Statistical Analysis

Experiment-specific details regarding sample size and statistical test used can be found in the corresponding Figure Legends. Significant p-values under 0.05 are denoted on all graphs and p-values above 0.05 are considered non-significant (ns). All statistical analysis was performed in GraphPad Prism 9. All data points for each experiment are included (no outlier exclusion was performed). For all whole animal fluorescence analysis, a sample size of 24 animals was used, each animal is considered a biological replicate.

| REAGENT or RESOURCE | SOURCE | IDENTIFIER |
| --- | --- | --- |
| Antibodies |  |  |
| Bacterial and virus strains |  |  |
| E. coli Bacteria OP50 | Caenorhabditis Genetics Center(CGC) | RRID: WB-STRAIN: WBStrain00041969 |
| Biological samples |  |  |
| Chemicals, peptides, and recombinant proteins |  |  |
| Vitamin B12 | Sigma-Aldrich | V2876 |
| Ethyl methanesulfonate (EMS) | Sigma-Aldrich | M0880 |
| Qiazol | Qiagen | 79306 |
| Alt-R® S.p. Cas9 Nuclease V3 | IDT | 1081058 |
| Tetramethylrhodamine ethyl ester perchlorate (TMRE) | Sigma-Aldrich | 87917 |
| Critical commercial assays |  |  |
| RNeasy Mini Kit | Qiagen | 74104 |
| TURBO DNA-free Kit | ThermoFisher | AM1907 |
| Deposited data |  |  |
| Raw and analyzed RNA sequencing data:<br>Raw data available upon publication (NCBI GEO).<br>Analyzed data provided as a supporting document with submission. | This study |  |
| Experimental models: Cell lines |  |  |
| Experimental models: Organisms/strains |  |  |
| *C. elegans* wild isolate | CGC | RRID: WB-STRAIN: WBStrain00000001 |
| zcls13[*hsp-6p::GFP*] V | CGC | RRID: WB-STRAIN: WBStrain00034068 |
| *acdh-1(ok1489)* I | CGC | RRID: WB-STRAIN: WBStrain00036246 |
| wwls24[*acdh-1p::GFP* + *unc-119(+)*] | CGC | RRID: WB-STRAIN: WBStrain00040155 |
| *hphd-1(ok3580)* II | CGC | RRID: WB-STRAIN: WBStrain00033247 |
| *alh-8(ww48)* II | CGC | RRID: WB-STRAIN: WBStrain00040157 |
| *glp-1(e2144)* III | CGC | RRID: WB-STRAIN: WBStrain00007730 |
| *pcca-1(ok2282)* X | CGC | RRID: WB-STRAIN: WBStrain00032465 |
| *nuo-6(qm200)* I; zcls13[*hsp-6p::GFP*] V | Held et al eLife 2022 | MRP232 |
| *hoe-1(mpt14[HOE-1::GFP])* IV | Held et al eLife 2022 | MRP258 |
| *hoe-1(mpt67[ΔNES])/tmC25* IV; zcls13 V | Held et al eLife 2022 | MRP379 |
| mpt134;zcls13 V | This study | MRP644 |
| mpt135;zcls13 V | This study | MRP663 |
| mpt136;zcls13 V | This study | MRP646 |
| mpt137;zcls13 V | This study | MRP647 |
| mpt138;zcls13 V | This study | MRP666 |
| mpt140;zcls13 V | This study | MRP650 |
| *acdh-1(mpt142[G294E])*I; zcls13 V | This study | MRP797 |
| *acdh-1(mpt145[A365V])*I; zcls13 V | This study | MRP806 |
| *acdh-1(ok1489)* I; zcls13 V | This study | MRP800 |
| *acdh-1(mpt154[ΔMTS])*I; zcls13 V | This study | MRP845 |
| *acdh-1(mpt157[ΔMTS,G294E])*I; zcls13 V | This study | MRP834 |
| *hach-1(mpt152[III:5556870a>t])* III; zcls13 V | This study | MRP899 |
| *acdh-1(ok1489)* I; *hach-1(mpt152)* III; zcls13 V | This study | MRP846 |
| *hphd-1(ok3580)* II; zcls13 V | This study | MRP820 |
| *alh-8(ww48)* II; zcls13 V | This study | MRP819 |
| *glp-1(e2144)* III; zcls13 V | This study | MRP511 |
| mptSi1[*eft-3p::3XFLAG::hoe-1(isoB ORF)::unc-54* 3'UTR], *zcls13 V* | This study | MRP428 |
| *glp-1(e2144) III*; mptSi1, *zcls13 V* | This study | MRP525 |
| *acdh-1(ok1489) I*; *glp-1(e2144) III*; *zcls13 V* | This study | MRP812 |
| *acdh-1(ok1489) I*; *glp-1(e2144) III*; mptSi1, *zcls13V* | This study | MRP785 |
| *acdh-1(mpt154[ΔMTS]) I*; *glp-1(e2144) III*; *zcls13 V* | This study | MRP910 |
| *acdh-1(mpt154[ΔMTS]) I*; *glp-1(e2144) III*; mptSi1, *zcls13 V* | This study | MRP909 |
| *glp-1(e2144) III*; *zcls13 V*; *pcca-1(ok2282) X* | This study | MRP833 |
| *glp-1(e2144) III*; mptSi1, *zcls13 V*; *pcca-1(ok2282)X* | This study | MRP832 |
| *acdh-1(mpt161[ACDH-1::GFP])/tmC20 I* | This study | MRP854 |
| *glp-1(2144) III*; *wwls24* | This study | MRP807 |
| *ech-6(mpt188[ECH-6::GFP])/+ III* | This study | MRP929 |
| *glp-1(e2144)*, *ech-6(mpt188[ECH-6::GFP])/+ III* | This study | MRP969 |
| Oligonucleotides |  |  |
| A complete list can be found in Table S1 |  |  |
| Recombinant DNA |  |  |
| Plasmid: pJA245 | addgene | 21506 |
| Software and algorithms |  |  |
| ImageJ/Fiji |  |  |
| GraphPad Prism |  |  |
| Integrative Genomics Viewer (IGV) |  |  |
| Trimmomatic-0.39 |  |  |
| STAR 2.5.4 |  |  |
| Salmon |  |  |
| Tximport |  |  |
| DeSeq2 |  |  |
| BioRender |  |  |
| Adobe Illustrator |  |  |
| Other |  |  |

