## Supplemental_Figures for "Germline status and micronutrient availability regulate a somatic mitochondrial quality control pathway via short-chain fatty acid metabolism"

#### Figure Supplemental 1

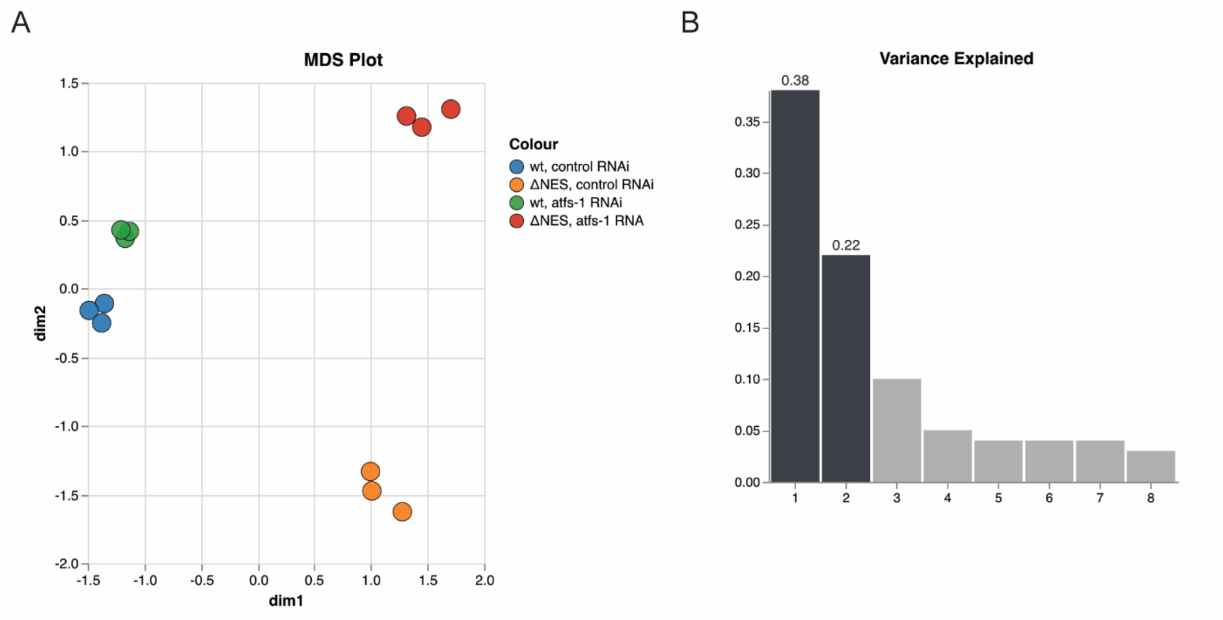


**Figure Supplemental 1: Increased nuclear activity of HOE-1 has a strong mitochondrial signature.**

GlimmaMDS was used to generate a multidimensional scaling (MDS) plot based on the data to visualize variation between RNA sequencing samples in the DESeq2 matrix of gene expression. (A) The two dimensions that account for the greatest variance between the RNA sequencing samples. (B) The eigenvalues of each dimension.

### Figure Supplemental 2

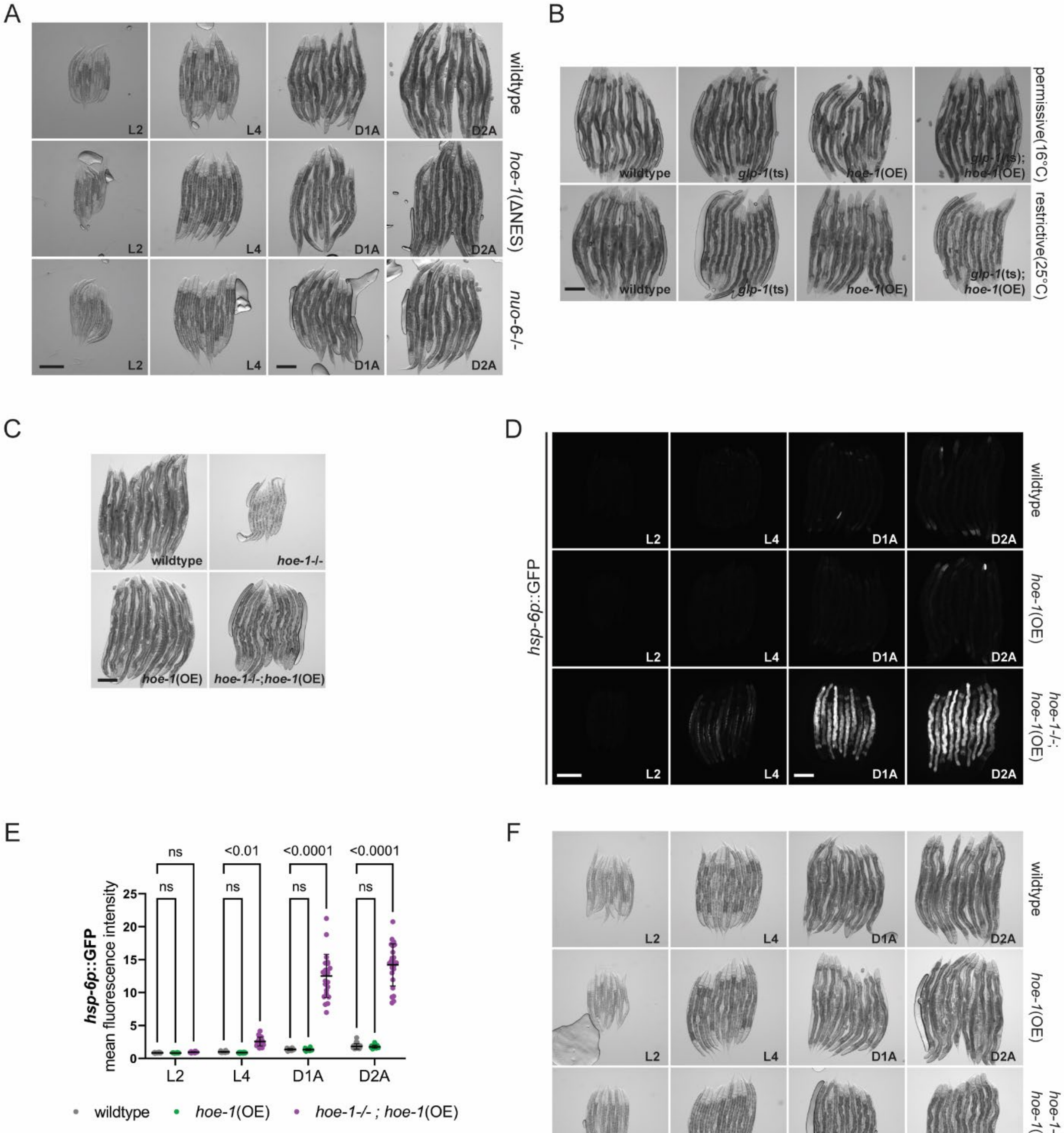

**Figure Supplemental 2: HOE-1-dependent UPR$^{mt}$ has distinct features.**

(A) Corresponding brightfield images to Figure 2C. Scale Bar 200 µm – L2 and L4 share scale bar, D1A and D2A share scale bar. (B) Corresponding brightfield images to Figure 2E. Scale Bar 200 µm. (C) Brightfield images of day 2 adult wildtype, *hoe-1* loss-of-function (*hoe-1-/-*), *hoe-1* somatic overexpression (*hoe-1*(OE), and *hoe-1-/-*;*hoe-1*(OE) animals. Scale bar 200 µm. (D) Fluorescence images of UPR$^{mt}$ reporter (*hsp-6p*::GFP) activation in wildtype, *hoe-1*(OE), and *hoe-1-/-*;*hoe-1*(OE) animals across development: larval stage 2 (L2), larval stage 4 (L4), day 1 adult (D1A), day 2 adult (D2A). Scale bar 200 µm – L2 and L4 share scale bar, D1A and D2A share scale bar. (E) Relative mean fluorescence intensity quantification of *hsp-6p*::GFP in wildtype, *hoe-1*(OE), and *hoe-1-/-*;*hoe-1*(OE) double-mutant animals across development normalized to wildtype L4 animals (n=24, mean and SD shown, ordinary two-way ANOVA with Tukey's multiple comparisons test). (F) Corresponding brightfield images to Figure S2D. Scale Bar 200 µm.

### Figure Supplemental 3

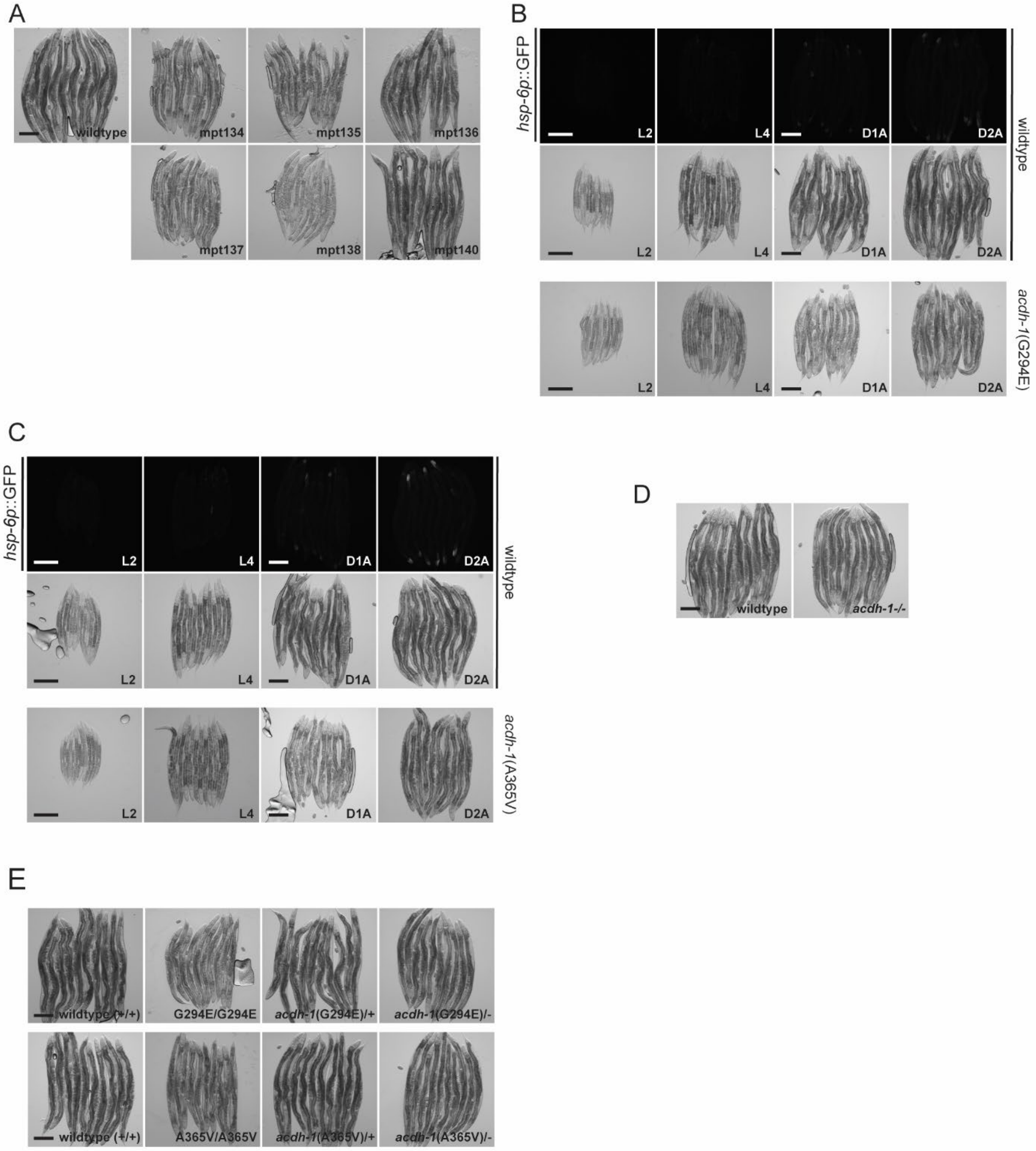

**Figure Supplemental 3: Forward genetic screen reveals gain-of-function acdh-1 mutants induce post-developmental, intestinal specific UPR$^{mt}$.**

(A) Corresponding brightfield images to Figure 3B as well as wildtype animals for comparison. Scale Bar 200 µm. (B) Fluorescence images of UPR$^{mt}$ reporter (*hsp-6p*::GFP) activation in wildtype control animals corresponding to *acdh-1*(G294E) in Figure 3D across development: larval stage 2 (L2), larval stage 4 (L4), day 1 adult (D1A), day 2 adult (D2A). Corresponding brightfield images for these wildtype animals and the *acdh-1*(G294E) in Figure 3D provided as well. Scale bar 200 µm – L2 and L4 share scale bar, D1A and D2A share scale bar. (C) Fluorescence images of UPR$^{mt}$ reporter (*hsp-6p*::GFP) activation in wildtype control animals corresponding to *acdh-1*(A365V) in Figure 3D across development: larval stage 2 (L2), larval stage 4 (L4), day 1 adult (D1A), day 2 adult (D2A). Corresponding brightfield images for these wildtype animals and the *acdh-1*(A365V) in Figure 3D provided as well. Scale bar 200 µm – L2 and L4 share scale bar, D1A and D2A share scale bar. (D) Corresponding brightfield images to Figure 3G. Scale Bar 200 µm. (E) Corresponding brightfield images to Figure 3I. Scale Bar 200 µm.

#### Figure Supplemental 4

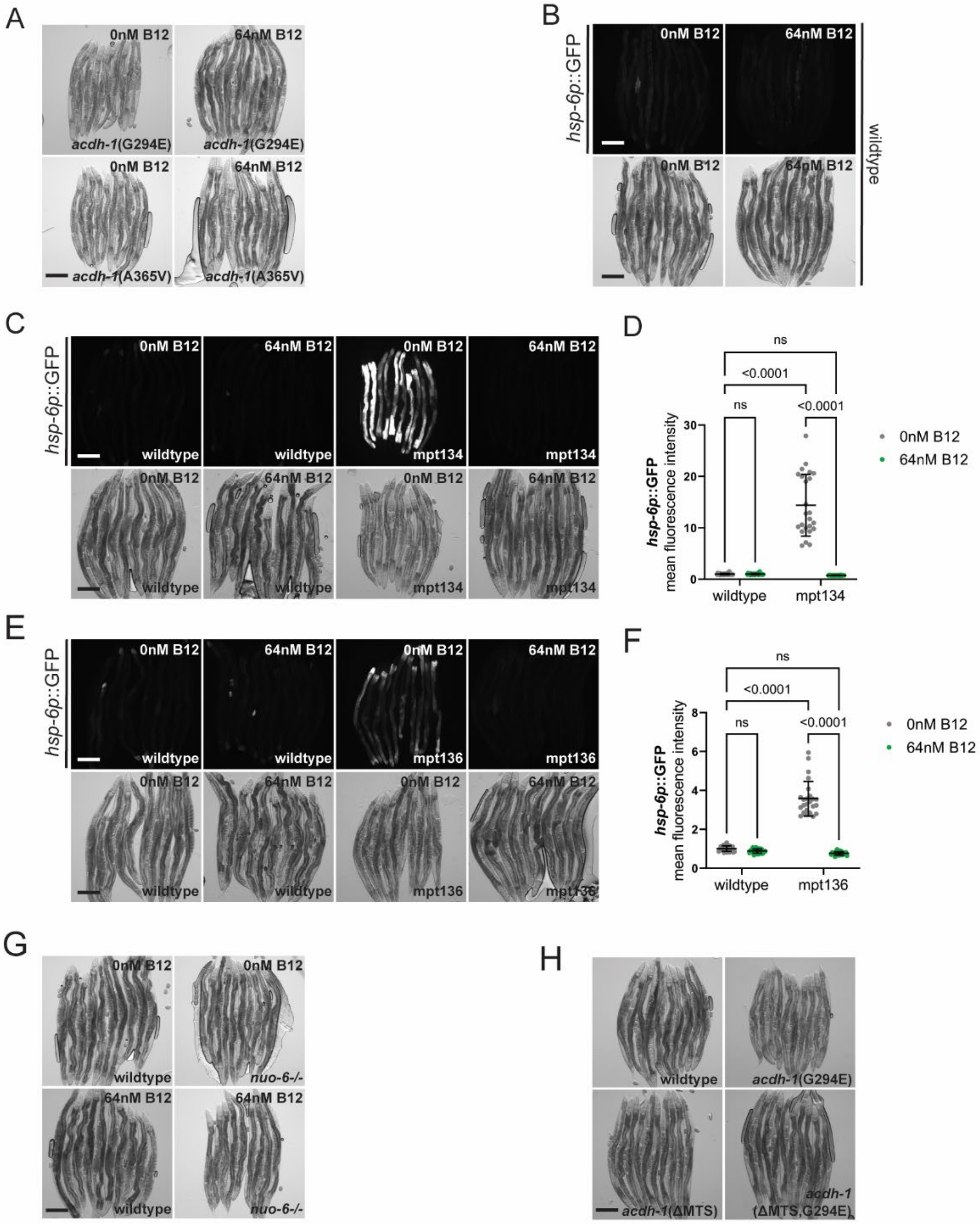

**Figure Supplemental 4: ACDH-1 is dependent upon vitamin B12 and required in the mitochondria to activate UPR$^{mt}$**

(A) Corresponding brightfield images to Figure 4B. Scale Bar 200 µm. (B) Fluorescence images of UPR$^{mt}$ reporter (*hsp-6p*::GFP) activation in wildtype control animals associated with Figure 4B on 0 and 64 nM vitamin B12 supplementation. Corresponding brightfield images for these wildtype animals are provided as well. Scale bar 200 µm. (C) Fluorescence images of UPR$^{mt}$ reporter (*hsp-6p*::GFP) activation in wildtype and mpt134 animals on 0 or 64 nM vitamin B12 supplementation. Corresponding brightfield images provided as well. Scale bar 200 µm. (D) Relative mean fluorescence intensity quantification of *hsp-6p*::GFP in wildtype and mpt134 animals on 0 or 64 nM vitamin B12 supplementation (n=24, mean and SD shown, ordinary two-way ANOVA with Tukey's multiple comparisons test). (E) Fluorescence images of UPR$^{mt}$ reporter (*hsp-6p*::GFP) activation in wildtype and mpt136 animals on 0 or 64 nM vitamin B12 supplementation. (F) Relative mean fluorescence intensity quantification of *hsp-6p*::GFP in wildtype and mpt136 animals on 0 or 64 nM vitamin B12 supplementation (n=24, mean and SD shown, ordinary two-way ANOVA with Tukey's multiple comparisons test). (G) Corresponding brightfield images to Figure 4D. Scale Bar 200 µm. (H) Corresponding brightfield images to Figure 4G. Scale Bar 200 µm.

#### Figure Supplemental 5

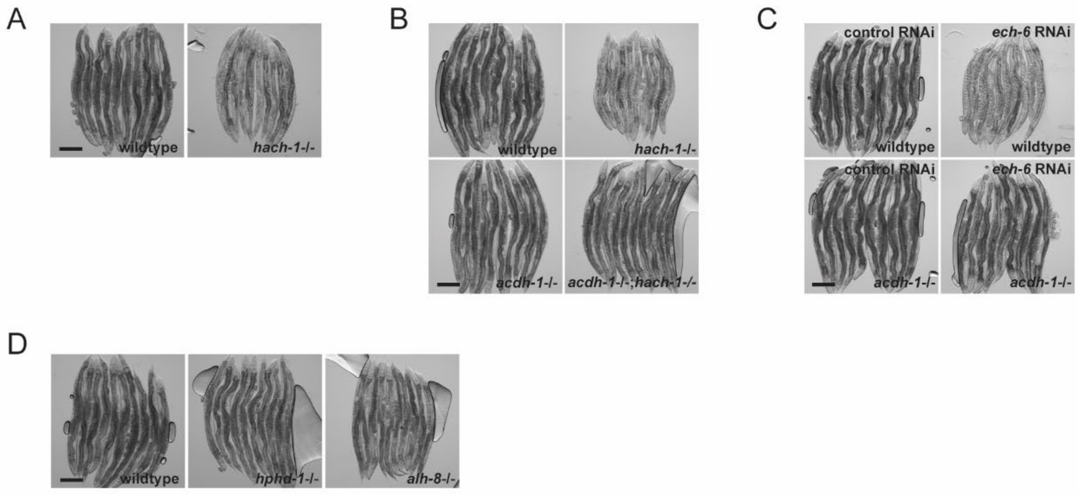


**Figure Supplemental 5: Acrylyl-CoA, an intermediate metabolite downstream of ACDH-1 and upstream of ECH-6, likely signals UPRmt activation.**
(A) Corresponding brightfield images to Figure 5A. Scale Bar 200 µm. (B) Corresponding brightfield images to Figure 5C. Scale Bar 200 µm. (C) Corresponding brightfield images to Figure 5E. Scale Bar 200 µm. (D) Corresponding brightfield images to Figure 5G. Scale Bar 200 µm.

#### Figure Supplemental 6

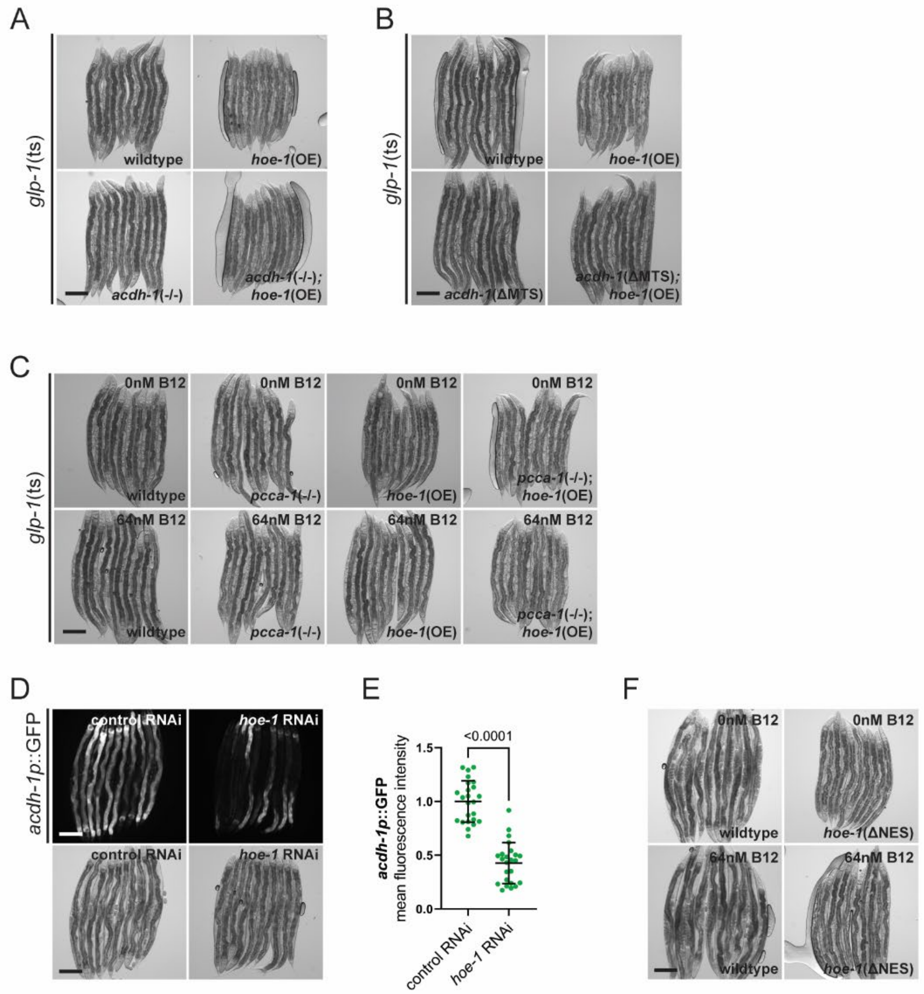


**Figure Supplemental 6: HOE-1 functions downstream of ACDH-1**

(A) Corresponding brightfield images to Figure 6A. Scale Bar 200 µm. (B) Corresponding brightfield images to Figure 6C. Scale Bar 200 µm. (C) Corresponding brightfield images to Figure 6E. Scale Bar 200 µm. (D) Fluorescence images of *acdh-1* transcriptional reporter (*acdh-1p*::GFP) activation in day 2 adult wildtype animals on control and *hoe-1* RNAi. Corresponding brightfield images provided as well. Scale Bar 200 µm. (E) Relative mean fluorescence intensity quantification of *acdh-1p*::GFP in day 2 adult wildtype animals on control and *hoe-1* RNAi (n=24, mean and SD shown, unpaired t-test). (F) Corresponding brightfield images to Figure 6G. Scale Bar 200 µm.

**Figure Supplemental 7**

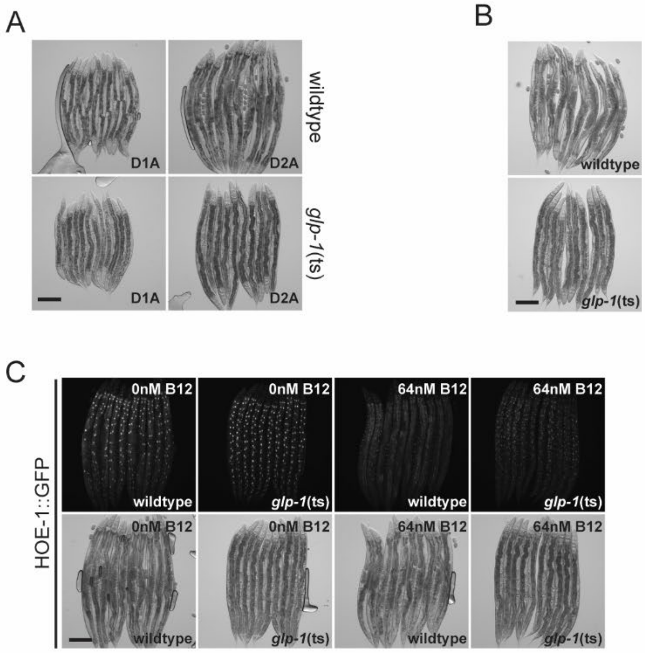


**Figure Supplemental 7: The germline non-cell autonomously regulates ACDH-1 pathway activity, influencing nuclear accumulation of HOE-1**

(A) Corresponding brightfield images to Figure 7A. Scale Bar 200 µm. (B) Corresponding brightfield images to Figure 7C. Scale Bar 200 µm. (C) Full worm fluorescence images of HOE-1::GFP corresponding to zoomed in images in Figure 7E. Corresponding brightfield images provided as well. Scale bar 200 µm.
